## Supplementary Table 2 for "Characterization of 475 novel, putative small RNAs (sRNAs) in Carbon-starved *Salmonella enterica* serovar typhimurium"

| **Supplemental Table 2. Novel intergenic small RNAs in Carbon-starved *Salmonella* enterica.** | | | | | |
| --- | --- | --- | --- | --- | --- |
| sRNA | Sequence | Log^a^ | 5h^a^ | 24h^a^ | Putative Gene Targets^b^ |
| sRNA63452 | TTCGGCATCACCGGAACCTGAACCAGTTCATACTTACTTTTCTTTCAGAAAAAATTTGGATGAAAATGGGATACGACATGCGTATCCCATCCA | 9 | 11 | 7 | - |
| sRNA93879 | CTTTATGCGCATCCATTTATTCCCATGCGAATAAACGGATGCCTGAACAGGCAGGGACGCCGGAAAA | 4 | 44 | 6 | CBW16188 |
| sRNA128997 | TACAGAATATAATGGCAACCGGGTGATATTGCATAACCTGTAGGCCCGGTAAG | 103 | 49 | 1 | - |
| sRNA157877 | ATTCGACTGGTTAACCTGGCACTCTCTCCGGCCAGGCGAGCCA | 357 | 125 | 143 | CBW18713 |
| sRNA212387 | TGCCCGATGGCGCTGCGCTTATCAGGCCTACATCCGTTGAGGATCTGTAGGCCGGATAAGACGCGTCGCGTCGCCATCCGGCACGGTCAGAAA | 15 | 3 | 2 | CBW20304 |
|  |  |  |  |  | CBW16297 |
|  |  |  |  |  | CBW16297 |
|  |  |  |  |  | SL1344_0073 |
|  |  |  |  |  | CBW20284 |
|  |  |  |  |  | CBW20331 |
|  |  |  |  |  | CBW16189 |
| sRNA215306 | TTACTCTATGTCTTTCGCATTACGGCAGACGGGCGTAAGAGAGCAACGCGACCTGGGCTGTAAGCATCATTAACGCCGCAGCAATG | 20 | 10 | 1 | - |
| sRNA237149 | AAAAAACCGGGTGGCGGCTTCGCCTTACCCGGCCTACGTTGTTTCTGGACGTTCTCTTTTACAGTTTCGCAAACACCCGACGCGCGGCGTCAATAGTATT | 51 | 27 | 27 | CBW16305 |
|  |  |  |  |  | CBW16305 |
|  |  |  |  |  | CBW16297 |
| sRNA248024 | TTTTCTATTATACGGCCTGGCAAGCGCGCTGGTATTTGGCCCGCTTTTTTTTC | 29 | 17 | 2 | CBW16124 |
| sRNA253928 | TTTGAGAATCATTCTGAACTTCGCCTTTCGTGCCACAGGTGCGTTGGCGACATTTTTTC | 23 | 48 | 12 | - |
| sRNA257035 | ATTGACGGGTCTTATTACCTGGCAGAAATTAAACGAGACTATACTTAGCACATCTTTATATTGTGTGACCGTCTGGTCT | 89 | 91 | 4 | - |
| sRNA266462 | GTCACATTCATACGATTGCGGCAGGCCGTGTTAT | 33 | 7 | 34 | - |
| sRNA289055 | TGTGGGCACTCGAAGATACGGATTCTTAACGTCCTCGGACGAAAAATGA | 3973 | 583 | 21 | - |
| sRNA351062 | CAAACTGGCTGCGCCAATAACGCCTGGTGGGATAGGCTCT | 67 | 686 | 48 | CBW18578 |
| sRNA361912 | AAGTCTCCTTGGCGGTGGTCTGAAAAACGTTCTTACAGGCCACAGGGTTAGCGGAAT | 284 | 32 | 3 | CBW19031 |
| sRNA437524 | TGCTATCGGGTGAGTCCTGAAAAACACTCACCCGAAATTTCATACTTCCTCCATGCGCTCTGTTTCTATAATTTGGGAA | 49 | 6 | 1 | - |
| sRNA441095 | ACGCATGAACCACTCGGAAGGGATTCTGAGTACCCCA | 1625 | 3652 | 27 | - |
| sRNA450983 | CAGTAGAACACAGGTTTCTGGCGGAGAAGGCTTTCCA | 13 | 10 | 0 | - |
| sRNA455513 | TTTTCTCTGCCTTATCCGGTCTGCAGGCAACGGTAGACCGGATAA | 80 | 7 | 0 | - |
| sRNA466560 | ATACATTGTCACTGGAATCCCGATCGCAAGGTTGGGATTTTTTTG | 29 | 2 | 0 | - |
| sRNA471433 | GTCACAGGCCGGAAAGATAGCGGGCGCTCGATACCCGCCGTCTCCA | 130 | 319 | 14 | - |
| sRNA485637 | GTCGCTTATTCGGCCTACAGACTGTACCGTTTTGTAGGCCGGATAAGGTGTTTACGCCGCCATC | 177 | 1041 | 652 | CBW20304 |
|  |  |  |  |  | CBW16297 |
|  |  |  |  |  | CBW18887 |
| sRNA487181 | TCCTCCTCACCTGCAACAGCGCCAGGTGTTATGGCGCTAATTATAGCGTCATCAGCTAAGCTTCTCGTTAAGCCTCGTT | 37 | 24 | 24 | CBW19913 |
| sRNA494586 | TCAGGTAGACCCAAAATCCGAAGATTTTGTTCTGGCCTGCGTCATGGTGCCCGTGTTCGTGCGCGTGGGCAGTCGCGTGCGTTAAAGTAT | 166 | 14 | 2 | - |
| sRNA501027 | CTTCGGGATGATTCTGACGACAGGGAATGTGATTGATTACGAGAACATCCCGGTTCCGCGAAGCAAAT | 538 | 86 | 3 | - |
| sRNA504956 | ATGTTATAGGTGGTATGGAGCACAGCTATACTATCTGATTACCTGGCGGACACTAAACTAAGAGAGAGCTCTATGAATCCTGAGC | 75 | 73 | 53 | - |
| sRNA507874 | CTCGGGGGATGTGACAAAGTACAAGGGCGCATCAACTGATGTGCCTTTTTT | 28 | 6 | 0 | - |
| sRNA543297 | TTTCTGAACAAACGGAGGCGGCATTCGCGGCCTCCGCTTGAAATCA | 90 | 10 | 13 | CBW19713 |
|  |  |  |  |  | CBW19339 |
| sRNA546177 | TTTCGCGTAGTGAAGCACTTATCAGGCCTGCTGTTATCCCGCAAGCCTGATAAGCGTAACGCCATCAGGCAATCGGTGCCGGATGGCGGCCT | 253 | 68 | 0 | CBW20304 |
|  |  |  |  |  | CBW20331 |
| sRNA561827 | TAGCTAAACCTCCTGTTGCGTTGTCCATAAACCGATTTTGCTGACGAATT | 155 | 12 | 1 | - |
| sRNA622126 | TATGCCGCGATTCGTCGGGTGGCGCTTGCGCCATCCGACAATGTGT | 17 | 1 | 1 | - |
| sRNA654423 | GCGTTTATCCGACTGACCAAATCGGTCTGGTCAGTCGGATAAGACTCCGATA | 100 | 12 | 7 | CBW16860 |
| sRNA655601 | ATCATTCAAGAAGCATCGCAACGGCAAGGGAAGAAATCCCCGCGGGCATAGATAACTGTGTGACCGGGGTTTCTGATC | 147 | 3 | 0 | - |
| sRNA740234 | GCTTATCAGACCTACAATGTTGTTCGAACCGTAGGCCTGAAAGCTATGCGTCATCAGGCAGAGGACAGGTTTAGATAACGT | 30 | 21 | 0 | - |
| sRNA748884 | TTTTATTTTTTCGCTGTTCGCTTTTGTCGGCAGCAATTTATACGTCAAAGAGGATTAACA | 27 | 30 | 2 | - |
| sRNA755730 | ATCAACATCAAGCGGCAGGAAAGAGGAGGATATAAAAAAGCCAACCGGGCGGTTGGCTCTTCG | 26 | 18 | 0 | - |
| sRNA773787 | AGGTAACGAGGTTCAATTTTTCGGTGCTATTCACCGATTCAATCTATACCGGGGCGGGTAGCGTCGGTATAAAGGGGGGGACGTAT | 48 | 5 | 11 | - |
| sRNA800767 | AATAATCGAGATGTTCAGAGCGAGACAAGGCGCGGTCTGAAGGAATCTTCGGGAGCATAGTTAACTATGTGACCGGAGTGAGCGAAGGCCGCAACGCAGT | 123 | 192 | 250 | SL1344_4445A |
| sRNA811833 | TTGCATCGCGGCGGCAAGGCTGCGAATCCCCAGGAGCTTACATCAGTAAGTGACTGGGGTGTGCAGCCGCAGCCAACAAAA | 48 | 27 | 29 | CBW20496 |
| sRNA844835 | GTTTAATCGTTAATGGGTATGAATAACCGCTGGGAAACCCCTGA | 30 | 0 | 0 | - |
| sRNA865865 | TTTAATCACAACTGAAGAAAATTAGCGTTGAGGCTGAACCCAACGCTAATTTTCCCTCTC | 16 | 22 | 53 | - |
| sRNA869761 | AACCACAGTAAACACTCTAGCCTCTGCACCTGGGTCATTTTTGATACGGTGCTTTGGCCGT | 94 | 145 | 27 | - |
| sRNA880025 | TCATAATGCCGGATGGCGGCGTGAACGCCTTATCCGGCCTACGACGACAATTAT | 17 | 5 | 0 | CBW20304 |
|  |  |  |  |  | CBW20304 |
|  |  |  |  |  | CBW18887 |
|  |  |  |  |  | CBW16297 |
|  |  |  |  |  | CBW16297 |
|  |  |  |  |  | CBW20552 |
| sRNA886206 | ATTTTATCCGGCAACATGCCCGGTAGCGCGACGCTTACCGGGCCTACGGACT | 122 | 4 | 0 | CBW18173 |
|  |  |  |  |  | CBW18086 |
| sRNA895467 | GAGCTGTAGGCCCGGTGAGCGTCCTCACTACCGG | 306 | 86 | 10 | - |
| sRNA898557 | AAATTGATGGATAAATAGACTGTTAGTTTAGGAATAATCTGGATGAATGTATCGCGCCGCAC | 75 | 103 | 129 | - |
| sRNA908052 | AGTAAAGGGATTGAAAGCTAAACACTGACGCCCGGTGGCGTTGTGTTTACCGGGCCTACGTATCCTGGCGCGTAGGTCCGGTAAGACGGGTGC | 24 | 16 | 12 | - |
| sRNA935960 | TAATAACTCCGTAGCCGGGTGCGCGTCGCATC | 43 | 37 | 3 | - |
| sRNA961825 | ATTATATCCTTTCAATGATAGCGTATCAGCTTGATAATGCGTTTGAGATCGTGTGCTTAGCTAACCCGGGAGATTCACTATTATATCCTTTCAATGATAGCGTATCAGCTTGATAATGCGTTTGAGATCGTGTGCTTAGCTAACCCGGGAGATTCACTATGCAGTTTTCAACAACCCCA | 62 | 36 | 36 | CBW16965 |
| sRNA993011 | GGAAGAAAAAACTGTGTTATGTATGTGCTGCATAATCATGCATGTAAATACCATGTTTACCGGGCTAGTGAAATCTACGCATGGCGTGGACAGACGCCAT | 315 | 77 | 28 | - |
| sRNA993137 | CAACGGTCTTCTCACCATAGACCAGGCATTGCGCGCCGTTAATCCCTCTGGGTTTCGGTCTATCGTGATGGGCAGCGACTCTGAAAA | 98 | 68 | 2 | - |
| sRNA993819 | AATCCATACAGCAACAGTGCCGGGGCAACCCGGTGCTGTTGTCCGTTTTAGCATCGGGCGGGAAAAGCCTGGAACCTGGAGAGCCTTTTT | 115 | 92 | 37 | - |
| sRNA1000128 | AAATATGATGCTCCAGGTATCATGGCGTTGATGATGAATCTCGTTATGCCTGATAGCACGTTGCTTATGAGGTCCGCGGGTATAGCGCAATGGATGCGTT | 41 | 19 | 0 | - |
| sRNA1015188 | TAATAAAACGGGGCCGCAAAGGCGCCCCGTAATATAACGCAGCCGAGAGGGTAAACCTACCCTAAG | 25 | 4 | 5 | - |
| sRNA1021440 | ATTATTCCGCATCTGATTACAGACAAAACTGGTTTTTGCACGCAACGTTAACGGTCGCGAGCGTTGGCGC | 2 | 17 | 1 | - |
| sRNA1024018 | CTAAAGGCGAGTAATCCTCTGAAACTTCGGGTCTTTTAGTCCGAAGTATAACGAGAT | 55 | 6 | 10 | - |
| sRNA1024473 | TTTTAACGCAAAACTTGAACCCGAAAAAAGCACCGTCAGGGTGCTTTTTTCA | 2 | 20 | 2 | - |
| sRNA1043460 | ATGTTTGGCTTCAACGGGGCGTTTCGCTGTAGACCCCCGTTGTGGCTGGGGGAGGGCCATCTTGA | 36 | 60 | 61 | - |
| sRNA1044917 | ATGAAAAAGCCTGCACATTGCAGGCTTTTTTATCGGTCACGCCAGTCGGCAGCTTTTTACAGTA | 53 | 12 | 1 | - |
| sRNA1047489 | AATAAATTCCGTCAATAAAACTTTACAGAAATATTGATGAGAGTTTGGTGTCTTTATGTGTCT | 23 | 40 | 0 | - |
| sRNA1048006 | AAAAAGGCCAGCATAGCTGGCCTTGAGTGTGACGTAAAAA | 43 | 3 | 8 | - |
| sRNA1061964 | AGCCGTTCTCATCCAGAACTTGCTGGATACGCTTGGCGAAAATGGGATGAAGATTTTTGTTCAT | 69 | 26 | 47 | CBW20435 |
| sRNA1066222 | AGAGGGAGTCCTATCAGATCTTGCTGATAATTTGCGGGTGACT | 26 | 6 | 2 | - |
| sRNA1116549 | TCACGCCATGCAAGTTTCGTAATGGTCAAAGTTGAGGTTTTTTAGTCTGTTGTAAAA | 29 | 7 | 0 | - |
| sRNA1127014 | CACCTCCCGGGAAATAGATACCCTTTATATGAGGATACTATTCCACTTTTCAAGTTACATGACGCGACAGGCAAAC | 35 | 21 | 25 | CBW17116 |
| sRNA1131800 | ATACTGTTAGAATATTTCCCGAAACAACGGACTACGCGAGTCTTTAGTTTCTTTT | 23 | 54 | 17 | - |
| sRNA1141227 | GGTCCTGGGCAGTGCTCGCCAAAACGCGTTAGCGTTTTGAACGCCGCTTGCGGCGGCCCGAAGGGCGAGCGTAGCGAGTCAAA | 34 | 154 | 70 | SL1344_3673A |
|  |  |  |  |  | SL1344_0083A |
| sRNA1180363 | ATTGAAAGCCGGATGGTGGTGTTATTGGCCGGTGGCGCTGCGCTTATGTAGACCGGATAAGGCGCCCGTGAACTGTGCCGACATCCGGCAATGGACGGGG | 18 | 12 | 16 | CBW20304 |
|  |  |  |  |  | CBW16297 |
| sRNA1189081 | ATAAAAAGAGCGACTCTGTTGAAAGCCCTGCTGTACACTTTGCAGATAAGGTGAGACAAAGGGGGGGGG | 23 | 0 | 0 | - |
| sRNA1198603 | AGCGTTCCACAGTATGATGCGCGGTTTGCGCTATCATCAATTTAATTATGAGCGTACT | 21 | 89 | 209 | - |
| sRNA1208607 | ACTGGCGTGCTGAAGACGTTACAGGAAGGAGTAGGTATAGAATGTTTGGCTATCGTAGTAACGT | 16 | 8 | 1 | CBW17200 |
| sRNA1210988 | ATAATGGCAAACCGTGGGGAGTCTGTAAGGCCTGATAAGACGTTTTAACGTCGCCGTCAGGCGCGA | 25 | 3 | 0 | - |
| sRNA1227939 | CCCGCAGCCCGCTAACTGTCTGAAAAATCAATACGTCTTACGCCATTGCTGCGTTGATGATCGGTCAGACAAAAT | 46 | 8 | 1 | - |
| sRNA1242409 | AAATAAAAGGGCACTTAGATGTCCTGTCCACGGCGGGGTTCTCCCCCCTCGCCAATGCGTGAGAACGTAGAAAAGCACAAATACTCAGGAGCACA | 201 | 14 | 12 | - |
| sRNA1243941 | CTAAGCAGTAATGATGTTGGGGAGAATTAAGGCAGCCATTCGGCTGCCTTTTTTAAA | 88 | 6 | 2 | - |
| sRNA1286671 | ATCGCACGTGGAAGTAAAGGTCGCTAACCTGTTGTCGTGTCAAGAGGTTAGCGGCACTCTC | 39 | 15 | 1 | - |
| sRNA1287943 | AAAAATAATGCCAGCTATATTTGATGGCAATCAAGATTTCCGGGTGGCTAAAAGTAGTCGTTCGGCACCATTGTAGATAAA | 2 | 13 | 2 | - |
| sRNA1291754 | TCAGTTGGCAACGGGTGTCATATCTTCAGGTATGGCGCCCGGAGC | 22 | 78 | 291 | - |
| sRNA1320514 | TAGCACTGCGATCTTGCCGACAAATGGTAAGAAAATCCTGCAGTGTGAACTCTTCGTCCTTGGCAGCTTCGTAAC | 18 | 37 | 10 | CBW19943 |
| sRNA1320716 | AAAAAAAGCAACTCTCATGCCATTTTATTGGTCAGAAGATTAACGTGTCCGGTGATACACCTTCTT | 2 | 116 | 295 | - |
| sRNA1321782 | ATTCTGATTGATTGAATTTGCCGGAAAAACCACGACAAAGCCTGCGTTATGCGGGCTTTGTG | 273 | 105 | 16 | - |
| sRNA1324246 | AAGCGCCGCCATCCGGCAGGTCATGCCTGATGGCGGCTGACGCCTTTCAGATCCTATAGTAACGCCGTTAT | 67 | 46 | 5 | CBW19985 |
| sRNA1327889 | CGTGACGTATCCGGCCCAAAGATCTATGCAATGGTCTCACCGTTGCGTGATGTATTGACCGGATACGATACCTGCTCCGCTA | 31 | 14 | 2 | - |
| sRNA1335449 | AGACCGAATAAGCGTAAAGCTATCTGGCCTGACGACGGGAT | 36 | 2 | 0 | CBW17747 |
| sRNA1337711 | TTTTTTTATCATTGTCGTTAAGGTCGTTATTCCTGCCGGGAAGCGTCGCGCTATCCCGGCCTTACG | 93 | 1 | 0 | CBW17758 |
| sRNA1356394 | TTTTTTCTGATTTTTAGCCTTAGAGCCTATCCC | 37 | 209 | 2 | - |
| sRNA1356450 | TTTGGACACGGACAGCGCGCAAAAACCGGAGCGTACACGTAGTACGTGAGGATTTTGAGCACTGCCCAGGACA | 101 | 259 | 52 | CBW17725 |
|  |  |  |  |  | CBW20596 |
| sRNA1366344 | GCCGGACGAAGATAGCGATTTTCGTCTGTGTCGAAGGTTGTGCGCCAATTTAGCAATGGTTGGCTAGATGGATAC | 14 | 10 | 4 | - |
| sRNA1368901 | GCCGGACGAAAATAGCGATTTTCGTCTGTGTCGAAGGTTGTGCGCCAATTTAGCAATGGTTGGCTAGATGGATTC | 19 | 16 | 3 | - |
| sRNA1372484 | ATTACGTCCAATGAACCGGTGTCTTCACCGACACAATGAGGAAAACCATGTCACATCA | 107 | 11 | 0 | - |
| sRNA1407102 | ATTAATAAAGTCTGTGTAGGCCGGATAAGGCGTTAGCCGCCATCCGGCTATGTA | 47 | 45 | 367 | CBW20304 |
|  |  |  |  |  | CBW16297 |
|  |  |  |  |  | CBW18887 |
| sRNA1414696 | ACAGCGTTCTACCTGATGTCAAATCCTTTGTCGGACGGCGACG | 1 | 14 | 5 | - |
| sRNA1418228 | TCTGGCTTCGGGCGCGCACGGCCCGGAAGCTAAGCCTGTGTTGCTAAAGTGATGATGCCTC | 26 | 30 | 99 | - |
| sRNA1468453 | ACTCAGGAGCCGGACGCAGAGTAATCTGTTCAAGGAGAAAGCGGCCTGTAAAGGTCGCTTTTTT | 220 | 88 | 9 | CBW17461 |
| sRNA1483192 | TTTCATCGGTAACTCCTTTTTGCTGAGGTTTAAAGTATGGTAGTGCGGGGACGAGCAAAAAAAAAGCCGCTAACGCACAGTTAGCGGCCTGAATG | 170 | 88 | 29 | CBW18736 |
| sRNA1485358 | TCCAGGCCGAAGGTGGGAGCGCGCTGGCGTGACGGACGCACTCCGACCATATCAAAT | 69 | 8 | 0 | - |
| sRNA1494275 | ATGTCGCATTTCTCCGCCTGGGATAGCATTAAACCGCAGGCGGAGAGAGGGC | 39 | 7 | 4 | CBW16219 |
| sRNA1497574 | TAAAAATGAAACCAGGCGGATTAAACCGCCTGGAGGAGTTTTACGTCT | 10 | 22 | 5 | - |
| sRNA1498969 | ACTCGCTTTGAGTCGGGCAATGGAACACCTGCCCAGAGAATAACCATACCGAGCGGTAAGTGAGAGCACAATGTCAAACAAACCCTTCTT | 25 | 129 | 24 | CBW17494 |
|  |  |  |  |  | CBW17296 |
| sRNA1500722 | TTTTGCCGGAGAACGGCGCAAGCGCCTTATCCGGTCTACACCGTTGTGTTGTTATCGTTAATACTCATAAGACCGCAAACATGTGAGCAATGTCGACACA | 17 | 37 | 2 | CBW20304 |
| sRNA1515557 | TCAGAAACCCCGGTCACATCGTTATCTATGCTCCCGGGGATTTCTTCCCTTGCCGCCGCGATGCAT | 37 | 23 | 41 | SL1344_4445A |
| sRNA1588729 | CTCCGGAACCACCACGCAACGATTTCCCAGTGTAGTGGTTCTTATCATGTGTATAT | 18 | 5 | 2 | - |
| sRNA1600950 | ACAATAGCCGCAGACGTGGTCTACGGCGCGTTGAGGTGATAGAAATACGGCAGGATCTCAGATAGACCGGGAGCGAGCCCTCCGTCTGAAA | 8 | 59 | 3 | - |
| sRNA1648615 | TAAGCACTATCAGTATTGGCCTTCTGCCCTACCGCTAAACATCTCATTGTTGTT | 1 | 19 | 4 | - |
| sRNA1676281 | ATGTCAGTCTTGTAAGAACGTAACATCACAACTGCGATACACCGCCGACGGGTAGCGAAAATATCTCAACCCT | 7 | 26 | 1 | - |
| sRNA1706658 | ACAGGGTCAGTGGGAATCGGGTCCGACAAGGACGAGTGTAGATATAATAAGGTCAAAAGACAGCTCATATTACCCTCCAGAATGTCAT | 4 | 58 | 21 | - |
| sRNA1723671 | TTATCTTCACTATACAGGGAACAACCAAATAACTGACAACACAAATCAAGCCGAGACCCGGAATTCGGCTATAC | 31 | 8 | 2 | - |
| sRNA1732903 | GAGCCGGATTACGACGCCGCGCTTAATGTCCTGAAGGCATAAGCCAGCGCCTGCTGGCGCTACGTTTGTCAGTGCTACAGGCGGTGC | 38 | 11 | 3 | CBW17707 |
|  |  |  |  |  | CBW17707 |
| sRNA1753147 | ACTCAACGGCCTTTGCTGAGAAGGCAAAGGCCGTTTCTTTTCTTCTTCTCCCTTCCGTCATGCTCTAAGTTTAAT | 91 | 210 | 151 | - |
| sRNA1755373 | GTTAATGGCGACGATAATAGGGAGAACAGCCTGTGCAGACAGGCATAGC | 142 | 16 | 0 | - |
| sRNA1759471 | TTTCTGCGGCCCATATACCACCCATAGGCAGCGGCTACAGGCAACAACAGAAATAACAACTCCAACATAAAT | 178 | 74 | 2 | - |
| sRNA1764707 | TGCCCGCTGCTGCGGTGTGGGCGTCCAGTGTTGCGGAACAGACCGAAAAACGGTCGTATCCGGGGTGT | 49 | 29 | 135 | - |
| sRNA1765147 | GCGGGACGGGTAATTCAACACTATTTTGCTGATCATTACCCGTAATACTTCGAGGGAT | 17 | 4 | 0 | - |
| sRNA1769418 | ACGCGCTATAGCACGACGGAATTTTGGTCGAATGTCAAGCAATTCTGTTGCCAGATTGACGAAA | 14 | 38 | 42 | - |
| sRNA1775237 | AAAGTTCACATGAAGAGGGTATCTAAAATGGCAGCGACATTTGCATTACACGGTTGGTGGCGCACTTCCTGATAGCGGGCGGTGTATGAAC | 5 | 54 | 0 | - |
| sRNA1805711 | AAAATACTTATAAATCTGCCGCAGATAGTAACACTCGTCGGGAAAGGCCGGTAAAGCAATTTCCGCTCACTCTTCC | 10 | 55 | 63 | - |
| sRNA1834529 | ACACCGCCTGGATAGGATTTTGCCTGGCCCGCACAGTTTTCGGCAGATTCTTTC | 45 | 6 | 0 | - |
| sRNA1858392 | ATAAGGTCGCCTGGCAACGTCGACCCGCCCCCAAGTGGACCGGTCGACGCCTTAGCC | 158 | 33 | 3 | - |
| sRNA1867048 | ACGGTTGGTTATTCATGGGGTAACTCCTCAGGATAGGCAGCGATATACTGC | 0 | 19 | 17 | - |
| sRNA1875281 | CTCTATTCCACCCCACAGCATAAAGTATGGGGGGTAATAGTAGTGAGAGTCAT | 52 | 17 | 2 | - |
| sRNA1885987 | GAAAATCATGCGTTTTTCGTTGTAACCCTCATCTGAATCGATTCGCTTGCGGACGGCGATTCAAAAAAT | 25 | 9 | 9 | - |
| sRNA1894083 | GGAAATGGGGCGTATGGCACTGGCGCCATTTGCGCCCCGGTAAGCATTATCGTTCTGCAACCAGACGAATGATGTCCTCATCCTCTTTTGGT | 159 | 15 | 5 | CBW17862 |
|  |  |  |  |  | - |
| sRNA1912451 | TGACTGTCCGCCCTGTGCCAGAAGCGGACGCTGATCATGGTGTGTACAGGCCAACAATACGATAT | 49 | 40 | 4 | CBW18613 |
| sRNA1913155 | GCCACTGGTTAGAAATCAGTAATACGATTTTATCAGGCTGGTGCTCTGGCCTGTCAGGATTTGAGGTATCCACCCGGTTTACCAGCCCCCTG | 7 | 16 | 4 | CBW18777 |
|  |  |  |  |  | CBW17085 |
|  |  |  |  |  | CBW18650 |
|  |  |  |  |  | CBW17894 |
| sRNA1915279 | TAATTGCCTGTCTCCGTTACTGGAGTGAACATATCAAAGACAGGCAATTACACAATTTTACTTACAGTCTCAGTTC | 10 | 19 | 11 | - |
| sRNA1918960 | ATTGGAATGGTTATGGGCTAACACGGAAGTTACAGT | 18 | 24 | 0 | - |
| sRNA1941471 | TTTTTGACCGTTGAGTAAAACTCCCCCACGCCTGTTCATCAGGTAGTACAGGG | 39 | 1 | 2 | CBW17916 |
| sRNA1942602 | CAAAAAAGCCGGCACACATCGCGTACCGGCTCTGTCAGCGCAT | 21 | 8 | 0 | CBW16545 |
| sRNA1969105 | GAGATAGCCGGATACGATAAGAAAGTCTCGTATCCGGCCGGGTTGACGGAT | 209 | 7 | 0 | - |
| sRNA1989185 | TGGCAAAGATTACCGGGCCCGGACGCCGTCCAGG | 37 | 24 | 1 | - |
| sRNA1990615 | ACAGATTACAGGGATACTCTCTTTATATCCACGCGCACGGTTGCCTTAATCATGTCTTCCGCGATAACACAGTTTGT | 17 | 14 | 0 | - |
| sRNA2001590 | AGACTGTGTCCCGCCATGGCGCATCGTCACGTCTCGCGAA | 30 | 13 | 15 | - |
| sRNA2009165 | ATTTTCCCGTCTCGTATGAAAATTCTTCCATACTCCAGAGGTCGGCTAAACGACTTCTGGAGCACGGAAGATA | 325 | 18 | 0 | - |
| sRNA2013506 | CGTCGCGTCCCCCGCCGCTGACGTAGGTAAGCGCCCGGGAGTTCGCTGCGGATGTGTTGTCAGCCCTGGCTTAAA | 17 | 0 | 0 | - |
| sRNA2020535 | GTAGCATGATTATCCGCGTCTTTTCCACGCTTTGTCGTGGACAGGACACGGGAT | 85 | 0 | 0 | CBW16452 |
| sRNA2037036 | GCAAAATGGCCTGAAGAAAATTATCAGAGAGAAAAAAACCTAAAGGAGATCTCAAGAGGAACAAATGATGAGAAATATTACAATCACTACTTCAGATAAG | 37 | 19 | 1 | - |
| sRNA2039748 | TGAGAGGGGGATTTGAACCCCCGGTAGAGTTGCCCCTACTCCGGTTTTCGAG | 62 | 38 | 9 | - |
| sRNA2073454 | TAGTCGATCCTCTGTAGTTAAGCACCCGTCTGGCGTGCAACCTTCGCCAGATACCGGGAGGCACCCGGCACCACAACGTT | 186 | 49 | 8 | CBW20037 |
| sRNA2145000 | ATTATCTTTACCCGGTTGATCTGATAAGGTTTGCCGGGTTTTTTT | 18 | 2 | 0 | CBW18085 |
|  |  |  |  |  | CBW20032 |
|  |  |  |  |  | CBW17024 |
| sRNA2176371 | CGCTACCGCCCCTGGCTCATCAGCTACCAGTGCACTGCGTACATATCGACTTGTTGCAAACCTCGCCCAGCAGGGCAAAGCTCACTAAAA | 141 | 114 | 17 | - |
| sRNA2202641 | GCCTGCTTACCAGGCAGGCCGGAGGCGCGTTAGCCTCGCCATCCGGCTCTGCCCTTTGGCCAAAAACTACAGCTTAT | 31 | 5 | 1 | - |
| sRNA2239569 | TCTCCTGCCCGCGTGAAGAAGGGACACGAGCAGTGTCAGACTTCCCTACGCTGGCATTATCCAGAT | 122 | 18 | 6 | - |
| sRNA2258848 | GCCAGACAGGGCGCTGCTATCGCGCCCTGTTGACGGCAGGCTTACAGTAG | 16 | 2 | 1 | - |
| sRNA2263199 | CGTGAAGCCCGGCGGCGCGATGCCTGCCGGGCCTGCGGCGACTTAA | 30 | 12 | 2 | CBW19290 |
|  |  |  |  |  | CBW16510 |
| sRNA2277763 | CTAAGCAAATGTGCCGGATAGCGACGCACTGGCGTCTTATCCGGCCTGACC | 28 | 9 | 1 | - |
| sRNA2288832 | TCGTGTTCTCCTGCAGGTGTGGCGCCTCTGCCCTGCATGGGGCAAAGTTTGTCAAT | 30 | 6 | 0 | - |
| sRNA2298229 | GGCGATGCCGCCCGGGCAGCCCTCCTCTTCACCTTTAT | 34 | 5 | 1 | - |
| sRNA2300307 | TAATCTTCATCAGGCAGCACCTCCCTTGGCTTACCT | 2 | 13 | 0 | CBW16138 |
| sRNA2311944 | TAATCTTCATCAGGCAGCACCTCCCTTGGCTTACCT | 591 | 554 | 2 | CBW16138 |
| sRNA2364092 | GTTAACCTGCGTTTCGCCTTTGAAGCCGATACGCATGTAGGTCTGGTCGCCATCGCTGCCTTTGTCGTCAGAGAAG | 31 | 7 | 5 | CBW18339 |
|  |  |  |  |  | CBW18339 |
|  |  |  |  |  | CBW17499 |
|  |  |  |  |  | CBW18018 |
|  |  |  |  |  | CBW17597 |
|  |  |  |  |  | CBW16417 |
|  |  |  |  |  | CBW18652 |
| sRNA2364248 | CGCCACCAGCAGAGCTGGTACCAGGAGGGACAGTACTTTAACTTTCATGTTATTA | 26 | 22 | 13 | - |
| sRNA2364333 | TATGCCACTGCTTACTGGTTAACCCTCATTAACCAGTCGGCAAGTCCATTCTCCGCAAAAA | 9 | 68 | 19 | - |
| sRNA2368373 | TTACCCAGCCTGATGGCGCTACGCTTATCAGGCCTACCAGTCGACTCATCTTTTACAGGCCGGACAGGCGACGCCGCCATCCGGCATTTTTTA | 80 | 86 | 20 | CBW20304 |
|  |  |  |  |  | CBW20331 |
|  |  |  |  |  | CBW16297 |
|  |  |  |  |  | CBW16297 |
|  |  |  |  |  | CBW18887 |
| sRNA2374312 | TTTCATGTGCGTTGGCTGCGAGTTACTCGGCCCATCCCTGGGCCTCG | 3 | 15 | 0 | SL1344_3673A |
|  |  |  |  |  | SL1344_4445A |
|  |  |  |  |  | SL1344_0083A |
| sRNA2381450 | TTTGTAGGCCGGATAAGGCGCTTGCGCCGCCATCC | 110 | 24 | 1 | CBW20304 |
|  |  |  |  |  | CBW16297 |
|  |  |  |  |  | CBW16297 |
| sRNA2385195 | ATGAGGCGTTTATGCCCTCATCCGGCGCGTCACT | 60 | 87 | 1 | - |
| sRNA2401404 | AGACAGTGCAGCTAACTTAATAGCAATACAATTAAAATGAAATTCCGCAACGGAAGACCAGGCCAGAAACATAAAAACAGCTTTTGGGC | 34 | 3 | 0 | - |
| sRNA2421910 | ACGTGTTGGGTTTCGGCTACGCCTCACCCAACGAGTCTATGGTCTCTATGCGTCGC | 267 | 17 | 1 | CBW20459 |
|  |  |  |  |  | CBW20080 |
| sRNA2436015 | GTCCGTGGTGGTCGGCAGTGCTTCGCGCTATTAAAATAGTGGTTAGCGAAGTGAAGTGCTGCTGCCGACAGGCGCAAAT | 121 | 29 | 1 | - |
| sRNA2439004 | AGGCGTCTACAGATTTCTGCTAATGATGGACGTGTAAA | 4 | 13 | 3 | - |
| sRNA2445415 | AGGTATCCTGTGTCGGCCTGTCAAAGGCCGGTAGTTTAAAATCATTGAGTCGTCAATATT | 15 | 8 | 7 | - |
| sRNA2457926 | ACAAGACGTAGGGAGAGAAAAAGTTCGCATCCGACCACAGCACAATAACAGGCTTACTCATAATGTCCTCTTGAGGGCGTAT | 32 | 14 | 11 | CBW18419 |
|  |  |  |  |  | CBW18419 |
|  |  |  |  |  | CBW20304 |
| sRNA2458808 | GTGTATACACGGAAAAAACCTGCGTCCGGCACCCTTATTCGTATTAAAAACCTGACATTAGGGAAGAGGAAATCCTCCCTAC | 18 | 16 | 15 | CBW20422 |
| sRNA2459810 | TATCCTGCCCGGTGAGCGATGCGCCGCCGGGCTTCTTTTATTTCAGCGAGCCTTTCAGGAAT | 39 | 13 | 7 | CBW18422 |
|  |  |  |  |  | CBW18459 |
| sRNA2462088 | TGCCGGATGGCGGCTGTGCCTTGCCCGGCCTACGATGCTCCCCGTAGGCCTGACACGCACAGCGCCATCAGGCAATGCAGA | 80 | 36 | 4 | CBW20304 |
|  |  |  |  |  | CBW16297 |
| sRNA2477353 | TTATTATAAGGGGCGCATAATGCCATTTTTGCCCCCAACAGACCATGAAT | 127 | 13 | 0 | CBW17380 |
| sRNA2495056 | GCCTGATCCAGCCTACGAACAGCCAGCGCTGTCGTAGGCCTGATAAGCGAAGCGCCATCAG | 112 | 162 | 17 | CBW20304 |
|  |  |  |  |  | CBW16297 |
| sRNA2495731 | TTTTACCTCAATGGCGTAAGTATAGTCAATCCTTGATTATTATTTCGCCACTAAGGAGGCATTCAGTGCGGATTCAT | 2 | 42 | 15 | - |
| sRNA2501436 | TCAAAAAGGTGAGCATTGCGCTCACCTTTTTATTTACTGCCGTTTATTCCGAGTCAATCTCTTTGAGTTCGTCCTGAAT | 30 | 18 | 29 | CBW18463 |
| sRNA2502279 | GACGGAACGCTACCGTATCCGCCTGTTTAGGTGTGAGCGATTTTAACAGCACGTCAAACGATGTCTATATGCGCGATCTCATT | 8 | 97 | 18 | - |
| sRNA2511779 | AAGGCTTCCATATAGCCTTACGGTCCCTCTCACTTC | 31 | 33 | 11 | - |
| sRNA2515627 | TACAGATGACACACTTTTGTAGGCCGGATAAG | 23 | 15 | 0 | CBW16297 |
| sRNA2542137 | ATGACGAGCGAAACGTCAGCGGCCATAAGTTGCCAGCAACGGATTGCGTAGAAC | 215 | 13 | 7 | - |
| sRNA2550290 | GTAAAAAAGCCCGGTATGCGTAGCGCCAC | 7 | 1 | 1 | - |
| sRNA2580004 | AAGTGACGCGGCCTGGTGCGCCGCACATTATCACCGTACTGCGCAACACAAA | 25 | 12 | 0 | CBW17383 |
| sRNA2584712 | TAGCTAAGGCGGTATCTTCCTTGCGCGCCTCAGGGTTCTCAAT | 86 | 34 | 0 | - |
| sRNA2594638 | AATAGCCTCTCTTATTATTATGGGTATTCTACGTAGTTAGCGGTATAGAGAGAAGTTCATTTAACCGATTGTTGCGATATCCTCTGGTTATGCTGTCTAT | 63 | 90 | 282 | - |
| sRNA2599691 | TATCACCTCATTCTCTCTTTGTTATACAGATGATCGTAGGCGGTCAGGCGCAGGATTACCACA | 20 | 0 | 0 | - |
| sRNA2639436 | GAGAAAAGTCCATTTCGTTGCAGTTCATTCCATGAAT | 11 | 22 | 0 | - |
| sRNA2650710 | AGGAAAACGGCCCCTACACCAGGAGCCGTTTTTTAAAACACACAACCGTGACGATTAACGCGTAAT | 19 | 3 | 1 | - |
| sRNA2656150 | AGGATTCAGGCGCCGACGAGCCGTAATGCCCGCCCACACCGCGAAAC | 81 | 15 | 4 | - |
| sRNA2657631 | AAGAAACGCCCCGATAAGCTTACTGCTCGTCCGGGGGCGCTGCATTGTACAAAT | 24 | 13 | 7 | - |
| sRNA2678384 | TCTGCGCGGGTTCCCCCAGTCATTTACTCAAGTAAGCGCCAGGGGATTCACTCGCTTGCCGCTTTTCTACAGCACGAATGCTCGGTAT | 394 | 407 | 56 | - |
| sRNA2681200 | ATTGCATAGCAGGGACAAGTCTGACATTCCCGAGTAATTTGGTCAACTATTTACTTGAGTAAATTAGTCAGGTATTTAGCGTTCCGTGC | 53 | 45 | 45 | - |
| sRNA2692792 | GATAAGGGTTTCCTGAACGAGAGTCTGACGAATTTCAACGGATTTCTTTTCAGCTTTGTGATGCAGATTTTTCACGTTGTTAC | 5 | 9 | 1 | CBW16275 |
| sRNA2694176 | ATCCTCTGAAAAAACGGGCGTTATCGTTAAGGCGAACGCCCGTTTTCCT | 175 | 10 | 86 | CBW18425 |
| sRNA2695893 | ATTTAACGCTGAACCATGACAAATAAAAGCGAAAAGATATTAAATTTTTCG | 31 | 20 | 0 | - |
| sRNA2723182 | AGCCCAGCTTCCGTTGAAGCCGATTAATTACGATGGGGTTCATGCGATACCAATAAATGAATGCGTCAACAATCCAGCACA | 40 | 19 | 0 | - |
| sRNA2726526 | CGCAGTCACGTTCTTGCAGACAACGTGACTGCGGTAATCCATCCCACCGGATTGTCTTCAAATTCTCCATGTTGCTGAATCGGCTAACAGCTTCTTAAAC | 7 | 20 | 16 | CBW17886 |
| sRNA2728577 | ATCAATAGCAAGGGCAAGGGCAAGGGCAAGGGCAAGGGC | 1 | 39 | 80 | CBW20290 |
| sRNA2763575 | GGTAAAACCCGCCCGTTACCGAGGAATCCCCAGCAATGGGTGGGTTGAATTGATGGGATTTTTTTAAGTTGCGATATCTTTGGCCTGACAAGGAT | 13 | 12 | 4 | - |
| sRNA2770084 | TCATTCAGGATTGAATATTCAAAATACAGTGGAAATGCAATAGCGGCTTTACTTACACGGCAACAATAATTTAACCCGGCCACTGTGCCGGGTTTTTTAA | 100 | 444 | 200 | - |
| sRNA2787726 | GTTTTATCGCCCGGATAACGGCGCGAAAGCGCCTTGTC | 31 | 21 | 0 | - |
| sRNA2806999 | ATTGTAGCTGGTCTTAACCGGGAGCAGGAACAGAGAATCTCCCGTAAAAC | 31 | 8 | 0 | - |
| sRNA2815179 | CTAATCATGCAACCATCTATGGCGTGGAGCAAACCCATAAAAAAGGGGCCAGCGTAATGCCAGCCCCTT | 7 | 23 | 8 | - |
| sRNA2829774 | TCGGTTGGTCGCCGCAGCGACACAGTTACAGAATTTT | 51 | 19 | 2 | - |
| sRNA2838573 | ACGAAAGCACTAGTCAGGCACAAAAAACAAAGGGTTATTCGGCGAGAAAACCAGACCTCCACCT | 1 | 149 | 391 | - |
| sRNA2893899 | AGTCTGATATATTGCTATACATCCAGAGATCACCCCAACAATATATCCAAAG | 19 | 3 | 0 | - |
| sRNA2922423 | AACTGCCGCAGTCCTGGGACTGCACCCGAGACAGCCACGACCTTAGAGGACTGTGCCTGGCTGCTATGTCAGGTTGGTCAAGCCC | 247 | 768 | 65 | - |
| sRNA2935133 | ATTAGGTCCCCTCCGGTAGCCCCCTAAGTTTTACTTTTCTCACGGAAT | 245 | 1 | 0 | - |
| sRNA2950623 | GCTGTGGCCGCGTTCCCTCACCCCAGTCACTTACTTTAGTAAGCTCCTGGGGATTCGCTCGCTTGCCGCCTTCCTGCAA | 10 | 38 | 30 | SL1344_4445A |
|  |  |  |  |  | CBW20496 |
|  |  |  |  |  | SL1344_0099A |
| sRNA2957079 | AACCGAACTGCTACATGTCGGCTCACATAATCACAAAAATCTTCTTCCGAACCTGAAAAATCCGGACGTCCGGCGT | 2 | 14 | 1 | - |
| sRNA2969723 | GAGCATGGCGCAACGTTCCCTGAGTCCCGGTTAATAGAGAC | 181 | 419 | 42 | - |
| sRNA2970828 | AGACAAAAGTGCTTTATGATTTCACTACCAACCATCACCTCATTCTGATCGATACGCTACAAGGAGCATTAACATGTTCTCTCCACAGTCACGCTTGCGT | 57 | 0 | 1 | CBW18882 |
| sRNA2981857 | TTTTTTATGCCGGATAAGGCGCAGTTCATGCGCCGCTAT | 207 | 4 | 1 | - |
| sRNA2984253 | ATTAATAACAGGAAAGCGGACAGGCCGGCACGCGTTACTGCTGGCCTTTCTG | 127 | 4 | 0 | CBW18875 |
|  |  |  |  |  | CBW18351 |
| sRNA2993440 | AGTGAGGTTGAAAAAAGTCATGAAGGGACAACGCGGGCGAGCGAGAAGAGAGGCTGCGTCTCACCCGAAAAGTGAGACGCGAAAACCTC | 198 | 36 | 77 | - |
| sRNA2993902 | CGTAAACGCCGCAGCACGGGCTGCGGCGCCAACGAACGCTTATAAT | 19 | 5 | 0 | CBW17288 |
| sRNA2996715 | GTCGCAGACGCTGTTCGCCGTTAAGCAACCCGCGTCCAGACGGAAAA | 24 | 11 | 0 | - |
| sRNA3026078 | ATACTGTAAGCCGGATAAGACGCATTTGCGTCGACATCCGGCATTAAC | 31 | 16 | 1 | CBW16297 |
| sRNA3032426 | AGTCTTATGGCGCTGGAAGGATTTCCTCTGGCAGGCAACCTT | 7 | 13 | 17 | CBW17623 |
| sRNA3072832 | ATAAAAAAACGAGTAAAATCAATATGAATGAGTCATTTGATAAGGACTTCTCCAAC | 19 | 13 | 4 | - |
| sRNA3114590 | TCGCGCCCCAGGAGCGTACATAAATACGTGACTGGGGTGAGTAAACGCAGTCAACTCAGATGCGGCTTGAAAGATGAAATGTATAAGCGATAAA | 16 | 66 | 0 | SL1344_4445A |
|  |  |  |  |  | CBW20496 |
| sRNA3116519 | ATTAAAACCGAACTCACCGGTTTATCCCCGCTGGCGCGGGGAACACTGC | 39 | 70 | 13 | - |
| sRNA3117025 | GGTTTATCCCCGCTAGCGCGGGGAACACAATCGCCAGCCTCGGAAA | 30 | 7 | 0 | CBW17603 |
| sRNA3130738 | ATATTCGCCGGATGGCGGCGTGACCGCCTTATC | 14 | 13 | 0 | CBW20304 |
|  |  |  |  |  | CBW18887 |
| sRNA3138538 | TGTTTCACCTTGTCTTACAAGACGCTTACAGACAAT | 39 | 12 | 0 | - |
| sRNA3162764 | ACAGTGCGGGCAGCATCCTACTTACCCGCGCAATAAACTCGCCGTCATCTCAGGTTAA | 21 | 15 | 12 | CBW17662 |
| sRNA3188055 | GTCAGTGCCCGGTGGCGCTACGCTTCCGGGCTTACAGAGACAATGTAAGCCCGATAACACGCACTCGCCGCCCGGCATAT | 16 | 5 | 2 | - |
| sRNA3217119 | ACCTGTAATCAACCTCTGGCGTTTTTCTGTATCCATCACAACCATTTCCGGTAAAGGTAAAAAAGGGATTTGAACCTTTCCGGAGATGATAAT | 21 | 62 | 15 | CBW20038 |
| sRNA3237780 | GGAAAGCCCCCCGGCGCGATAGCCGGGAAAGCAAAGTGTGACGGGTTTCAAA | 23 | 13 | 2 | CBW18874 |
| sRNA3239101 | GATAAGGCGAAGCCGCCATCCGGTATTTTTACGAACACGATGCCGGATGGCGGCGTGAACGCCTTATCCGGCCTACAGAT | 35 | 146 | 15 | CBW20304 |
|  |  |  |  |  | CBW20304 |
|  |  |  |  |  | CBW18887 |
|  |  |  |  |  | CBW16297 |
|  |  |  |  |  | CBW16297 |
|  |  |  |  |  | CBW16297 |
|  |  |  |  |  | SL1344_4369A |
| sRNA3241625 | GCATTTGCGCCAGTAATGCCTGGCGACGGCGTTGGTATTCCTGCTGAGTCATGACATGGCCTCCATTTCGTTTTTTCTTAGTGT | 27 | 11 | 3 | - |
| sRNA3242325 | TTTTGGACCCTGGTGAACCAGAAAAGGGCTTGTATCTTCACACCAGGGTAGCT | 31 | 9 | 1 | - |
| sRNA3243004 | TTCTGACTGGCGACAGCCATGACGCAACTTCCCGAACTCTCTTTATCACGACAAGAAATCCGCCGGATGATTCGGCAACGTCGCCGT | 53 | 102 | 405 | - |
| sRNA3245414 | AGGGCAAACGATTAAACGTGATACATGTCACCAAATTTGCCCTGACCGAATTTT | 197 | 192 | 1 | CBW18433 |
| sRNA3247244 | CGGCCTGCATGACCCATTATTCCCTCGTCACTTCTTTGAAGT | 4 | 25 | 0 | - |
| sRNA3249067 | GCTACGCTTATCCGCTCAATAACCAGAGAATACTGGAGGCGGATAAGCGCAGCGCCGGCAG | 94 | 221 | 21 | - |
| sRNA3250185 | AAAAAAATGGCTTCCCAAGGGAAGCCATTTCAATCAGCAGACTATATTACAGAACGTCAATCGCGTTCAGTTCTTTAAAT | 47 | 45 | 155 | CBW19143 |
|  |  |  |  |  | CBW19143 |
| sRNA3251305 | AGACATGTTTCGTTGTCCTGTATCGTCGGGCTTTGGGAAAAAA | 31 | 17 | 7 | - |
| sRNA3259138 | CCGTAACGACTTGACGACAGCGCGTTTTGGGCTACGCCGGAAAATTT | 16 | 4 | 0 | - |
| sRNA3260159 | GCGGCGTGAACGCCTTATCCGGCCTACAAGCATCAATCAGCTGTAGGCCGGATAAGCGCAGCGCCATCCGGCAAAACAA | 30 | 33 | 36 | CBW20304 |
|  |  |  |  |  | CBW20304 |
|  |  |  |  |  | SL1344_0073 |
|  |  |  |  |  | CBW16297 |
|  |  |  |  |  | CBW16297 |
|  |  |  |  |  | CBW18887 |
|  |  |  |  |  | CBW19132 |
| sRNA3274017 | ATTGCTAACCCAACCGGAGGGCATAATGCCTG | 8 | 12 | 3 | - |
| sRNA3278697 | AGGCAGCCGTCCTTTGGCAAAAGACGGCTTATATGAGTGACAACCGTCCGCTTTCCCCACATACTGAAC | 22 | 165 | 15 | - |
| sRNA3284953 | ACAAGGATAAGCCTGTCGCGCCAGCGGGTAAAATACCTGATGGCGCGACGCTTACC | 33 | 9 | 6 | CBW17842 |
| sRNA3290772 | TCGGCCTGTGAGTTATGCCGGATAAGCCCTTAGCGTCGCTA | 47 | 28 | 0 | - |
| sRNA3293424 | TGTTTCGCAGGCCGGATAAGGTGTAAATGCCTTAT | 23 | 4 | 0 | CBW16618 |
| sRNA3296804 | ATTTCGTTGACGAAGCGAGGCGATTACGGTTTAATGCGCCTCGTTGCCC | 75 | 19 | 1 | - |
| sRNA3307760 | GCAAGTGGGGAGCGCGTTATTCTCCCCAGTGATGCCAACGCTACTGACGAACAATTAA | 8 | 1 | 0 | CBW18335 |
| sRNA3312835 | TGCCCGGGTAATCCGTTACCCGACTGATATTAA | 2 | 15 | 11 | - |
| sRNA3327831 | GGTTTCGCCCCATAATCGATATATAAATAATTATATAACG | 8 | 2 | 0 | - |
| sRNA3339202 | TAAGGATCGTTCTGATTCTCCCTCTTACGGTAGTAAAAGTGTCTACGCTTAAAGGGTGTTCAACCTGAAGGGAGAAGCTCAATGTCTCATCT | 44 | 6 | 0 | - |
| sRNA3340225 | TAAAAAAGCCGGGTGGCGACTTCGCCTTACCCGGCCTACAGCGT | 35 | 6 | 7 | CBW16297 |
| sRNA3357552 | ATACCGTAGGCCGGATAAGACGCTACTGCGTCGTCATCCGGCAACGGCAGAGCGCCGGATGGCGGCTTAGATGCCTTGTCCGGCCTACGGGCGCCCCTGT | 35 | 12 | 7 | CBW16297 |
|  |  |  |  |  | CBW16297 |
|  |  |  |  |  | CBW20304 |
|  |  |  |  |  | CBW20304 |
|  |  |  |  |  | - |
| sRNA3364545 | ATTACCCTCGTCTGACGTCGCCCTGATGCGATGCGCAAGTGTCGCCATCAGGCTGCGGCGTCATGTCGGCT | 114 | 83 | 17 | - |
| sRNA3370753 | ATTCGCTGGCGCTGCGCTTATCAGACCTGTGCCCTTGTAGGTCTGGTAAGCGCGGCGCTACCTGTA | 95 | 267 | 10 | CBW16916 |
|  |  |  |  |  | CBW20304 |
| sRNA3393458 | ATGCCCGTCGTTTTGCTGTAAACCCGTTGGGGCGGCATGATGTCGCCCCCTTTCT | 407 | 1095 | 2762 | - |
| sRNA3394975 | ATCTTTTGAGAGCGGGCTGCTCTCAATTCAGATCGAGTTGTAGTTGTAAGGCCGTGCTTCCGAAAGGAATGCGCGGCTTGTTTTC | 377 | 245 | 36 | - |
| sRNA3410852 | ATTGCCTGCCGCCACGTCAAGGATGACGATCTGTAGAG | 19 | 43 | 0 | - |
| sRNA3460763 | AGCTTTACTCCCTGTTTCAATAGTCCTGTTATTTTGCCATATCCAATAAATCACATGGCGGCGTAAA | 28 | 5 | 2 | - |
| sRNA3466666 | ATGGTGCCGCCGATAATCACCACGCCGCCAAGCAGCGAAGGTGACGTTTGG | 107 | 154 | 3 | CBW16681 |
|  |  |  |  |  | CBW19540 |
|  |  |  |  |  | CBW20249 |
|  |  |  |  |  | CBW20088 |
|  |  |  |  |  | CBW16909 |
|  |  |  |  |  | CBW16264 |
| sRNA3469871 | GAACGTCCTGTTCCCCGGTGGCTCAACAAGGCGTCCAGCCAGGTCTAAAGCCCTCCGCACTGCGGAGGGCAAAAG | 57 | 28 | 0 | - |
| sRNA3472646 | TCAGCGATCCCAGTGTACGCCGCGTAAACTGTTAAATAATTTACGCGACGTTAAG | 287 | 10 | 0 | - |
| sRNA3489276 | GATAAAAGGCCGCTATGCGGCCTTTTTCCTTTCTTTACAATACACCAACAAT | 14 | 8 | 1 | CBW19119 |
| sRNA3493902 | TTTCCGGACCACAAAAAACTTACTCAGCAACGATGCTGATGTATTTACGGTTTTTCGGGCCTTTAACTTCGAAT | 70 | 7 | 0 | CBW19371 |
|  |  |  |  |  | CBW19371 |
| sRNA3498005 | GCAACTTATCCGGCCTACGGGGATCATGCGAAGTGTAGGCCGGATAAACCATCCGGCAT | 31 | 53 | 45 | - |
| sRNA3515728 | CCGCTGGAAGCTTTCTGGATGAGCCGTCTG | 52 | 27 | 0 | - |
| sRNA3515800 | ATTTACCAATGCGCAAAGGACACAAAGAGGGCGAATGCGAGGTAAGCGTATGATGCGCCAGCCCCGTCGCCATGCTCTTGCTGTGCCCGCGCGCCGTCGG | 219 | 10 | 0 | CBW17017 |
| sRNA3521831 | CTTCCGGCTCCCCGCTTACTGCCTTGGCTGATGGCTGGG | 466 | 46 | 4 | - |
| sRNA3534732 | TGACAAAGTCCGGTAAACCTACCCCCTCGGATATGTTTGGCCGGCACAGAGAAAGTTTGGGAAAAAACTCTCTTGTAACGTGGGGT | 184 | 21 | 0 | - |
| sRNA3549132 | ATGCGGAGTCGCGCAATTTTGGCTTTTATGCATTTAATTTGCGTAATCCGGATCGGCTGCAACCAGAGAGCAACGTTCTACAGGT | 13 | 57 | 25 | - |
| sRNA3564889 | GCGTCGGAGAGGCAGGACAATCCAGGGATAATCCTGCCCGTATCCAACTACTCTTCGCTGAACAGGTCGCCGCCGTGCAT | 121 | 23 | 2 | CBW19442 |
|  |  |  |  |  | CBW19442 |
|  |  |  |  |  | CBW20211 |
| sRNA3593555 | TGTGGGCACTCGAAGATACGGATTCTTAACGTCCTAGGACGAAAAATGAAA | 7140 | 1007 | 32 | - |
| sRNA3620977 | TCCTTACTATAGCGCTCTCCCCTTCAGGAGAGAGCACGGGAT | 55 | 6 | 1 | - |
| sRNA3623172 | TTCTTTAGGCGTTCAGTTAAGTTATCCAAAACGGGCGACTTACCTGAAGCGCCCGCCTGGTGA | 229 | 18 | 2 | - |
| sRNA3623731 | CCGTTGTTAATTCAGGATTGTCCAAAACTCTACGAGTTTAGTTTGACATTAATTTAAAACGTTTGGCCTTACTTAACGGAGAA | 276 | 47 | 3 | - |
| sRNA3624232 | AATAAAACGCCCTCACACAATATGAGGACGCCGAATTTTAGGGCGGTGTCGAAAAGGTGTCAAGAA | 6 | 42 | 16 | - |
| sRNA3627609 | AGACGAAAAAAAGCGGGGTTAACCCGCTTTTTTTGTGCCCGATGATTTATCGAACCAATGATAAT | 21 | 11 | 0 | - |
| sRNA3627758 | GCGTAGCCAACACCGCGGTGGCTTGAAAGACGACGGGGAT | 24 | 7 | 0 | - |
| sRNA3681038 | GCGTAGCCAACACCGCGGTGGCTTGAAAGACGACGGGGAT | 14 | 14 | 5 | - |
| sRNA3685306 | GTTGATTATTTACTCTTTCCTTCGTTGGCTAAACATCGGGTCTCCTGCCGCCCCCCTGAGCGCCGCATGAGGTATACATCCAGTTAGTAAGAAACAAGTA | 215 | 66 | 13 | - |
| sRNA3691557 | CGATGGCGCTTGCGCCTGTCGGGCCTTGTCAG | 8 | 20 | 15 | - |
| sRNA3693224 | TTTAAGCGAGGCGGCTTGTAGGCCGGATAAGGCAGCAATGTGCGCCAGGATTGCCTGATGGCGCTGCGCTTATCAGGCCTACGTGCTCTC | 24 | 54 | 5 | CBW20304 |
|  |  |  |  |  | CBW20304 |
|  |  |  |  |  | SL1344_0073 |
|  |  |  |  |  | CBW20331 |
|  |  |  |  |  | CBW16297 |
|  |  |  |  |  | CBW16916 |
| sRNA3711543 | CTTTCAATGGCGTCGTAATAAAATCCGGGGCGGTTTATACCGCCCTTTTCGA | 175 | 249 | 1052 | CBW19587 |
| sRNA3729870 | TCCCCGCCGGATGGCGGCGCTGACGCGCCTTATCCGGCCTACAGAAC | 22 | 6 | 1 | CBW20304 |
|  |  |  |  |  | CBW18887 |
|  |  |  |  |  | CBW16297 |
|  |  |  |  |  | CBW16297 |
|  |  |  |  |  | CBW19985 |
|  |  |  |  |  | CBW19668 |
|  |  |  |  |  | CBW16296 |
|  |  |  |  |  | CCF76853 |
| sRNA3755629 | TAAAGAAAGCCGGATGGCGCTGACGCTTATCCGGCCTACATGCCGGGAGAAAAGATAAA | 35 | 50 | 1 | SL1344_0073 |
|  |  |  |  |  | CBW20304 |
|  |  |  |  |  | CBW20304 |
|  |  |  |  |  | CBW16297 |
|  |  |  |  |  | CBW19097 |
| sRNA3758267 | ATGACGCGCGGGGAAGAGACAACAGGGACTCTTTCCCTGCGAACGGAAGCCCATTGCAGG | 56 | 29 | 53 | - |
| sRNA3773060 | AGACTAACGCCGGGGGCGCTTCGCTTGCCCGGCCTACGTTCGCACGTCTGTCGTTTGTAGGCCGGATGAGGCGCAGCCGCCATCCGGCAATAAA | 20 | 31 | 3 | CBW16297 |
|  |  |  |  |  | CBW20304 |
|  |  |  |  |  | CBW20304 |
|  |  |  |  |  | CBW18887 |
| sRNA3781793 | TCCTCCCCGGCGACCCGCCCACGGAGTTCCTCTCCGGGACTACCGCTCCCCGATACCCTGCCAGACAAT | 33 | 609 | 631 | - |
| sRNA3782057 | AGTTATGGTAAACGGCTGAGCTAGCCCCGCGCATAGAGTTCGCAGGACGCGGGTGACGCGGCGGCATAAGAAACGCCAGTAGCTCAATGGTCATCGACAA | 59 | 23 | 30 | - |
| sRNA3825805 | GAGAACGCGCATTTTTCATCTTTCCGTGACATCATTTATAATGTGTAAAAATGCAAAGCGCAGAGTTACAGGGCATCCTGCCGGGCAAA | 13 | 60 | 4 | - |
| sRNA3831578 | TGTGGCGCTACGCTTACCAGGCCTACAGGTACGGTGTAGGCTTGATAAGGCGCAGCCGCCATCCGGCATAAAAC | 164 | 59 | 493 | CBW20304 |
|  |  |  |  |  | CBW20304 |
|  |  |  |  |  | CBW16297 |
| sRNA3835685 | TTACGCCGGATGGCGCTGCGCTTATCCGGCCTACACAACACCGGGGCGCAGCGCTCTTTTAATTAT | 106 | 23 | 4 | CBW16297 |
|  |  |  |  |  | CBW20304 |
|  |  |  |  |  | CBW20304 |
|  |  |  |  |  | SL1344_0073 |
|  |  |  |  |  | CBW19132 |
| sRNA3855978 | TCTACAGGCCCGATAAGCGCTGGCTTGCGCTT | 49 | 12 | 0 | - |
| sRNA3856713 | AGAGCAGGACTTACCTGTCGCCATTGTTAATACTCCTTTCACCCAAAAAT | 21 | 4 | 0 | CBW19706 |
| sRNA3870656 | TCTACAGATGCCTTGTAAGCCGGATAGCGTAGCGCCATCCGGCACACAGAAT | 42 | 44 | 3 | CBW20331 |
|  |  |  |  |  | CBW16297 |
| sRNA3871532 | TCTACAGATGCCTTGTAAGCCGGATAGCGTAGCGCCATCCGGCACACAGAAT | 44 | 69 | 6 | CBW20331 |
|  |  |  |  |  | CBW16297 |
| sRNA3889188 | AGGTCTCCTGGTCGGATTGATACATTCCAACACCTTTTAT | 1 | 240 | 27 | - |
| sRNA3898508 | ATCCCCAAAATAGTTCGAGCTGCAGGAAGGCGGCAACGCAGTGAATCCCCAGGAGCGTACATAAGTACGTGACTGGGGTGAGC | 10 | 37 | 8 | SL1344_4445A |
|  |  |  |  |  | CBW20496 |
|  |  |  |  |  | SL1344_0099A |
| sRNA3901329 | CTTCCGCATCCCACGCTCTCAGGACACCCGTTGTACGGAGGATGTCTTGAGAGCATCGGGCTAT | 30 | 37 | 53 | - |
| sRNA3931763 | CTGGGAATAGGGTAGTGGGTCGAGACGAGTCCATTTCTGAAAAAGACGCTTATAAA | 33 | 1 | 0 | - |
| sRNA3940293 | CGCTACCGCCCCAGGCTTCACAGCAACCCGATGGCTAATACCGCACTCATTTTAGAT | 17 | 39 | 1 | CBW18626 |
| sRNA3986689 | GGTTCGGCAAGCAGCGCAGGAATAGCGTGCGTTAACTTGCCAGATGATTTACGTAAAAGTGACTGTCCAACATATTACT | 62 | 108 | 13 | - |
| sRNA4006485 | AATGTTTTCATGGTGCGCGCCTAAACATTTAGAGTTGTCTATAGACAATCATGTACAACTTC | 40 | 20 | 40 | - |
| sRNA4019564 | ATCGTAAACCGGCGCAACGAAGTCCTGGCTGAAACGGGTGGTGCCGTCAGCGCCTTAACCCC | 28 | 7 | 3 | - |
| sRNA4031693 | AGGCCAGGATTGCCAGAACACCCATCAGGCGTGCCCTTCTATTCACTAC | 2 | 17 | 21 | - |
| sRNA4066570 | AGAGATCTTCTGTTTCTCACAGATTCTTCCCTATTTATCCACAGGACTTTCCAGGAAAGGATAAGTGTAATCGATCCTGGGGAACTCCT | 8 | 82 | 34 | CBW19534 |
| sRNA4066734 | AGGGCGATCCTTGCGCTTTACCTGGCAGTCCGTATAAT | 1 | 57 | 1 | CBW20329 |
| sRNA4069379 | ATCCTGATTCGGTGTAGGCCGGATAAGGCGAAGCCGCCATCCGGCAAGGCGCGCCGGTCAGCCTGATGGCGCTAACGCTTATCAGGCCTACAA | 221 | 63 | 5 | CBW16297 |
|  |  |  |  |  | CBW16297 |
|  |  |  |  |  | CBW20304 |
|  |  |  |  |  | CBW20304 |
|  |  |  |  |  | SL1344_4369A |
|  |  |  |  |  | CBW20331 |
| sRNA4086010 | AGTTATATGTCTTGATTATGACGGTTTGATGACAATGGAAATAAAAAAAGCTGGCCCGGGGAGACACCAGACCAGCCTGCAGGGG | 21 | 3 | 1 | - |
| sRNA4090297 | AAAAAAAAGGCCGTCTGTCGACGGCCTTCTGCCTGGTACTACATTTGT | 373 | 17 | 3 | - |
| sRNA4092239 | TTCGGGCGCCCCGAGCCTTGTAGAGAGTGGGGTTATTTTTATAAGTACTGC | 81 | 1 | 0 | - |
| sRNA4110784 | ACATCAAGGCGGCAAAGGAGTGAATCCCCTGGCACTGACTTAAACGTGTGACTGGGGTAAGC | 21 | 5 | 0 | SL1344_4445A |
|  |  |  |  |  | SL1344_0099A |
| sRNA4133397 | ATAATTTGCGTCACAGGAAGGCGGTAACGCAGCGAATCCCCAGAAGCTTACTGTAGTAAGTGACTGGGGTGAGTGAGG | 26 | 1 | 0 | - |
| sRNA4140398 | TATACCTTGTCGGGGGGCGCTAGCGCTTACCTGACCTACAAAATCAAGCAGACGATGTAGGCCGGGTAAGGCG | 303 | 7 | 3 | - |
| sRNA4154553 | TTACTCTGGGAAAATACGTTCTTTGGCGTGAATAGCGGCATTGTGCGTATCGATGCCTCAGCGCCCGAATT | 15 | 2 | 0 | - |
| sRNA4162399 | GCACTGGACAGCGGGGTGTGAAGTCCGTATTATCCACCCCCGCAACG | 53 | 208 | 28 | - |
| sRNA4167628 | GCAGTGTCGTCAATGACAGTGAATAATGACGAGAAACCGCCAGCCCGT | 32 | 253 | 17 | CBW16638 |
| sRNA4167843 | TTTATCCCGAATGGCGACGGTCAAGTACTGACCTGCACCATGACGGGTAGCAACATCAGGCGATAC | 34 | 67 | 1 | - |
| sRNA4176959 | AGTTCTGTATAAATTCCCAGTTATTCTGGCGGATGGCGCTATGCTTATCCGCCCTACGATAAGTAGCCCCGGTAAGCGAAGCGCCACCGGGCGTTTTTTG | 42 | 27 | 6 | SL1344_0073 |
|  |  |  |  |  | CCF76769 |
| sRNA4179482 | ACGATTACGCTAAGAGGACATTCGCCTTGGACACACCCAGTAGATACTGGCTCATTATCCTGTCATCCAGGATCAACTCCTAAGGCTATCCCTTTTTGCT | 66 | 31 | 196 | - |
| sRNA4183113 | ATGGTTGGCAGAGAATGATTCAAGCTGAAAATACTTCGTCAGGAGAACGTCAAGAATGCGGGCGATTTTGAGAGGGTTGTTGCC | 103 | 36 | 6 | - |
| sRNA4191128 | GTAAACATCCAGACGTCTAAAACTCGTCATATTTCGACTTGCAGATGCGTTGGCT | 29 | 0 | 3 | CBW20496 |
| sRNA4195272 | GACGCTTTGTTGAGCCTACACCGACTCCGGTCTCGT | 26 | 77 | 0 | - |
| sRNA4206621 | ACTGAAAGCGGACATCGACATTTCAGGCTGCGGCGACTCAGCTTGAAAAATGTCGGTATT | 146 | 44 | 30 | - |
| sRNA4209588 | GCTCACCCCGGGCACATAGTTATCTATGTGCCCGGGGATTCGCTCAGTTGCCGCTGCGCTGCAAT | 9 | 68 | 20 | SL1344_4445A |
| sRNA4223041 | TAACTGTAAAACCGTAGGCCTGATAAGCGCAGCGCCATCAGGCAACCGGCACTGCCGGATGGCGGCTCCGCCTTATCCGG | 69 | 25 | 10 | CBW20304 |
|  |  |  |  |  | CBW20304 |
|  |  |  |  |  | CBW20304 |
|  |  |  |  |  | CBW16297 |
|  |  |  |  |  | SL1344_0073 |
|  |  |  |  |  | CBW20331 |
|  |  |  |  |  | SL1344_4369A |
|  |  |  |  |  | CBW16916 |
| sRNA4230634 | GCTCACACCGTAGGCCGGATAAGACGCGCAGCGGCGCCATCCGGCATAGCGGGTGATTATTCCCCGTCCTGAAT | 86 | 34 | 7 | CBW20304 |
|  |  |  |  |  | CBW16297 |
|  |  |  |  |  | CBW16297 |
|  |  |  |  |  | CBW20039 |
| sRNA4232507 | ATTGTCGCGGCGACAGAAATGGCTGCGCAGCCGGCTTGCCGCCCGTCATGGCGTGAAGCCGGTACGCCGTACCCGCCAGAGACGCCATCGGAAGGAGTGA | 87 | 89 | 79 | CBW18343 |
| sRNA4238588 | ATCGTGCAAAAGGGCTGCACCACGGTGACTATGTTGCACCAAAATAGTGCTTTAATGTGAACATTAAGCACCACATTGGTGCAACAGAACA | 23 | 1 | 0 | - |
| sRNA4240654 | CTTACGGATAAAAACCCTGCCGAAATGAAAGCTCGGCGGGGTTTTTTT | 9 | 0 | 0 | - |
| sRNA4269302 | CGCTCAGGGAGTGGTTGTTATTTTTCTTTGCGGACGGGCCCGCAAAGGTTCAGAATTGGATAAATTTTCCTCCATCCGGGAGGAGT | 18 | 70 | 23 | - |
| sRNA4288594 | ATAATTTGCATTGCACGTCTGTAGAAGCGAGTCTGATGACTCGCTTTTTTTTT | 183 | 121 | 3 | - |
| sRNA4294735 | TTTTGCGCGCCTGATGGCGCTACGCTTATCAGGCCTACAGGTTGCAGAAACGTGTAGGCCGGATTAGGCGTTAGCCGCCATCCGG | 34 | 27 | 46 | CBW20304 |
|  |  |  |  |  | CBW20331 |
|  |  |  |  |  | CBW16297 |
|  |  |  |  |  | CBW16297 |
|  |  |  |  |  | CBW18887 |
|  |  |  |  |  | SL1344_0073 |
| sRNA4311608 | ATAAATGCCGGATGGCGGCTTTACCTTATCAGGCCTACATAAGCACTCGGCTGGTAGGCCTGATAAGCGCAGCGCCATCAGACATTGATTGGCAATTAAG | 65 | 44 | 9 | CBW20304 |
|  |  |  |  |  | CBW20304 |
|  |  |  |  |  | CBW16297 |
|  |  |  |  |  | CBW16297 |
|  |  |  |  |  | SL1344_0073 |
|  |  |  |  |  | CBW16916 |
| sRNA4313724 | TAATTTGCCCGGTGGCGCTGACGCTTACCGGGCCTACAGATTGGTGATGATTTGCAGGCCGGATAAGCCCCAGCCGCCATTCGGTAAAACATCACCAGTA | 55 | 41 | 22 | CBW20304 |
|  |  |  |  |  | CBW16297 |
| sRNA4340853 | ATAACCTCCAGCTCCCTGCCGGGAGCTTTATTACAACTCATATTGATCTACATCTCTGT | 48 | 38 | 21 | - |
| sRNA4354661 | ATAGCGGCGCTTTGCGCCTTATCCGGACT | 17 | 1 | 0 | - |
| sRNA4359630 | GAGACGCCGGATGGCGCTGCGCTTATCCGGCCTACAAGACGGGGTAGGCCCGGTAAACATCACGTCACCGGGCAATAT | 126 | 51 | 3 | CBW16297 |
|  |  |  |  |  | CBW20304 |
|  |  |  |  |  | CBW20304 |
|  |  |  |  |  | SL1344_0073 |
|  |  |  |  |  | CBW19132 |
| sRNA4372453 | TGTGGGCACTCGAAGATACGGATTCTTAACGTCGCAAGACGAAAAATGAATACCAAGTCTCAAGAGTGAAC | 8269 | 1583 | 24 | - |
| sRNA4383182 | TTTCACACCAGAACCTGGCTCATCAGTGATTTTATTTGTCATAATCATTGCTGA | 167 | 49 | 0 | - |
| sRNA4403706 | TGTTGTAGACATAGCTCATTCCAAAAAATTAAGGACGTGGCTTGTCAGACGACGGATGGAGTAAT | 24 | 2 | 0 | - |
| sRNA4459705 | GCCATCCGGCAGCGTATTGCCTGATAGCGCTCACCCGCTGCCGTGTCAGGGCGATTATTCAAAT | 30 | 2 | 1 | - |
| sRNA4461150 | AAGGGCAGAACAGAAACCGGTGCAGGCGCAAATGCAAGTAAC | 24 | 2 | 0 | - |
| sRNA4463102 | ATGGCACTGCGCTTATCCGGTCTACGTAATGCATAGCGTCTGTAGGCCGGGTAAGGCGTTT | 17 | 31 | 0 | CBW20304 |
| sRNA4477374 | TGTCATGCCGGGCGGACCCGGCCTACAAAGCCAAAAT | 121 | 3 | 0 | - |
| sRNA4481167 | GTAAGCCGCCATCCGGCAATCGTGTAGCCTGAT | 33 | 4 | 0 | CBW17168 |
| sRNA4482999 | TTTCCCCTACCAGGAGTACATGGATGTGCTTCCCCC | 23 | 26 | 6 | - |
| sRNA4483054 | GCGTGCCCCTCTCTACTGCCGCCCGTTTTCCGT | 112 | 33 | 0 | - |
| sRNA4491788 | ATATTCAGGCCGGACGGCGATAACAGTGCCCTCCG | 17 | 3 | 1 | - |
| sRNA4498153 | TTCTCGTCTGCTGAAATGCCTGGTGTAAACCAGGCATTTTCTT | 24 | 13 | 43 | - |
| sRNA4527235 | AAGTTCAGTTGAAAAAGCGTTGATGATCGCTGGATAATCGTTTGCTTTTTTTTGCCACCCGTTTTGTATACGTGGAGCTAAACGTTTGCTTTTTTGCGGC | 1 | 18 | 5 | CBW17162 |
| sRNA4534828 | CCAGCCACATAGTGAACTTGAACTATGCTGCTGAGGGTGTGCTCGCTTGCCGCCTTCCTGTAACACGAAAT | 2 | 59 | 1 | CBW20496 |
|  |  |  |  |  | CBW19869 |
| sRNA4545622 | ATTGCCGGATGGCGCTTTGCTTATCCGGCCTACAGCGAAAGAGAATGTAGGCCGGATAAGGCGTTCA | 41 | 75 | 95 | CBW20304 |
|  |  |  |  |  | SL1344_0073 |
|  |  |  |  |  | CBW16297 |
|  |  |  |  |  | CBW16297 |
|  |  |  |  |  | CBW17168 |
| sRNA4545710 | TTCCGGGTTGCCTGATGGCGCTACGCTTATCAGGCCTACAACGACTACGTTCATCAGGCAATAAA | 56 | 26 | 2 | CBW20304 |
|  |  |  |  |  | CBW20331 |
|  |  |  |  |  | CBW16297 |
|  |  |  |  |  | SL1344_0073 |
| sRNA4550089 | ATTTTCAGGCAACCCTCCCGGTGATGCCAAAGAGAAAAGTGTAGTTCGTTGACAATAAAT | 36 | 11 | 0 | - |
| sRNA4551416 | CAGGCCCGGTAAGCAAGGCGCTACCGGGCAATGACACGGTAAATCAGTTCTTTTTCACGAAC | 38 | 13 | 0 | CBW20312 |
| sRNA4552158 | ATGTTTGGCTTCAACGGGGCGTTTCGCTGTAGACCCCCGTTGTGGCTGGGGGAGGGCCATCTTGA | 17 | 22 | 6 | - |
| sRNA4570153 | AGGCGTAGCACAAGCTCCATTGCTACAAACATAATTTTATTTAAGTGTCAGGAAAATTCCGGACAAATCCCTTTTTTT | 18 | 0 | 0 | - |
| sRNA4577680 | ATTGGTCGGGTTTTGATGGCGTGCCGTAATACTTGTGCCGCCATCGTGC | 8 | 19 | 0 | - |
| sRNA4591479 | AAAAGGCACGTCATCGTGACGTGCCTCTTTGGTACTACCCTGTACGATTACTGTTCGCT | 30 | 232 | 256 | - |
| sRNA4597370 | ATGCGTTGGCTAACCCCAGTCACTTACTTGAGTAAGTGACTGGGGATTAACA | 57 | 17 | 51 | - |
| sRNA4626825 | TCATTCAGGTCGTATTGAGGCGGTAGCTGAGAGAATCTCAGAAGCTCCCAACGAAGGGACCTGGGGTAAAAAAGCCGCCACTCAAGAC | 61 | 50 | 0 | CBW19735 |
|  |  |  |  |  | CBW18905 |
| sRNA4634066 | AAAAAGCGGCAGAGTGAGCAATACCCTCTTCAAACAGAAGAGGGTCAATT | 137 | 33 | 2 | CBW20387 |
| sRNA4697295 | ATTTGCCGTCGTTCAGGTTGCAGAAGTGTGGTTATTGTTCTGCAACCTGAA | 2 | 22 | 6 | - |
| sRNA4703739 | AGATGAAAATCGCCGGACAATAATACCCGTCATACTTCAGGCTGCAGGCGCGTTGGCTGCCTGCAACCCGAATTATTCAGGGTAT | 53 | 58 | 22 | CBW20455 |
| sRNA4713076 | GTGATGCCTGATGACGACGCGATTGCGTCTTATCAGGCCTACGGTCTTTGCAAACGGAAA | 45 | 16 | 4 | CBW16297 |
| sRNA4732259 | TTTTTGTGCCGGATGGCGGCGCATGCACGTTGCGATCTTGTAGGCCAACCGCCATCCGGCATCATCACGACATTAT | 403 | 306 | 137 | CBW19985 |
|  |  |  |  |  | CBW16433 |
| sRNA4740982 | TGATGGCGGCGCTAATGCGCCTTATCAGGCCTACGGGGTAAT | 140 | 33 | 8 | CBW20304 |
|  |  |  |  |  | CBW20304 |
| sRNA4742776 | ATTATTACGGCATTGGCACGCCAGAACAAGTTCTGAGAGGTGAATCCGCTGAGTATAATGATCTTAGCGATGATTTCGAC | 15 | 10 | 5 | - |
| sRNA4745126 | TAATCGTAGGCCTGATGGCGCTGCGCTTATCAGGCCTACCATGACCCGTAGGCCTGATAAGTCCTTTCCGCCGCCATCCGGCAGACC | 334 | 206 | 15 | CBW20304 |
|  |  |  |  |  | CBW20304 |
|  |  |  |  |  | CBW20304 |
| sRNA4756044 | GATAGGCGTCAGGTTTAGGGCAGGAGTGTGATTAGCCATCTTATCAGTGTTTCCCCAGGGTAAGATGGCTGTGA | 18 | 40 | 13 | CBW16297 |
| sRNA4772464 | AAAACAAACGATGATTGTAATGGGGGAAGGGGCGGGATGCC | 9 | 75 | 0 | SL1344_0073 |
| sRNA4776489 | GCATTGATAATCAGTCCGGCCTGAAAAGGTCGGGTAACTGATTATCAGATGATGACATTCTCCAGCATCAAAGCCTCGGGTTGAGTTGAAAGGTATTTAC | 614 | 621 | 96 | CBW20331 |
| sRNA4783501 | AGGGATAGTGATCCGTGCCGCCTTCAGGAAGGCAGCCACCAGCGGAGATGACGTTGCGTAGCGTTGGATCGTTTCGTGTTCAT | 395 | 129 | 12 | CBW169163 |
| sRNA4794013 | AGGAAGAGACGCGTTCTGAATCTCGTCAGAACGTGTGGCGATTACAGATAACAATCAGCACATCACACCTCGGGAAATTTACGTTGC | 18 | 32 | 20 | CCF76750 |
| sRNA4837089 | ATTCGTCCTCCCGACGTTTGTCGGGAGGCGTAATGTGCACCACACTAAAAATATCGCGAATGAGTAGCCTGAGCGCTCATAT | 11 | 103 | 67 | CBW20506 |
| sRNA4857626 | GCAATGCCTGATGCGACGCTTACCGCGTCTTATCAGGCCTACAGTTTCACAACGTACTGGAAT | 74 | 33 | 2 | - |
| sRNA4864124 | CCCATCCTTTCAACAACGAGCACCCGATAT | 49 | 16 | 6 | - |

^a^ Log –; 5h –; 24 --

^b^ Putative Gene Targets
