## Supplementary Table 3 for "Characterization of 475 novel, putative small RNAs (sRNAs) in Carbon-starved *Salmonella enterica* serovar typhimurium"

| **Supplemental Table 3. Novel intragenic small RNAs in Carbon-starved *Salmonella* enterica.** | | | | | | | |
| --- | --- | --- | --- | --- | --- | --- | --- |
| sRNA | Sequence | Log^b^ | 5h^b^ | | 24h^b^ | | Putative Gene Targets^c^ |
| sRNA1014845 | AGGGTTGTCAGCTTTCACACTAACTCTCTCTTTATTAAGTCGGCGACGAGATACTAAC | 17 | 9 | | 7 | | - |
| sRNA1065845 | ATACAGTTGTTGAATGTAGAGGGAATTATGAGAATTGAACTTGTTATCAGCCGGGCAAAACAGCTTCCGGAAGGTGCCGTTCC | 111 | 222 | | 1323 | | CBW17053 |
|  |  |  |  | |  | | CBW18685 |
| sRNA1125244 | GGTCGTGAGCAACCAACGCGATATGTTTGCGCGTCGGCAAGGTGCGAGTCGTCAGTTCCATAATTCGATTCCGTAGTTAACCT | 42 | 11 | | 4 | | CBW17112 |
| sRNA1161265 | CATATGCCCCCGCGCAGGCGGGGGCGTTTGTGTTATACGTTTTTACGTTCGATGATTTGTTCGCCCCAGAAGAGCGAGTCTTTGTCCGTTTTCTCAAAG | 18 | 49 | | 21 | | CBW17153 |
| sRNA1202532 | AATAAACAGCCCCTTTACGGGGCCGACAAATTATTGGCTAAAACGGGAAAGCCGGAACGGCGTCAGATCAAAG | 74 | 50 | | 35 | | CBW17193 |
| sRNA1205786 | GCGCGAGCGCGCCCTCCGAAATCAATTGGTGAAAGGAATAACGATTTCACCAGGTTTAACTT | 26 | 103 | | 164 | | CBW17197 |
| sRNA1276193 | CTCTACAACCAGTACGCGCATCATCTCTTCTCCCTTGTGTTAACAATAAGAACAGTCT | 47 | 44 | | 2 | | - |
| sRNA1308054 | TTTGCCGCCCGGCTGGCAACGCTCCAGTCTTAACCACCTGGACCCGTCGTCAACGGCGGGTCATTGC | 35 | 27 | | 2 | | CBW17300 |
| sRNA1416448 | ACCAGTTTAGTACGATTCATTATTAATACCCTCTAGATTGAGTTAATCTCCATGTAGC | 44 | 33 | | 32 | | - |
| sRNA1440139 | ATTTATTATCGTTACCATAACAGCGCACTTGTAGTGAGTGAGCAAGAGTAAAGTAAAAATATCTTAGAGCCTATCCC | 69 | 18 | | 13 | | CBW17426 |
| sRNA1476693 | AAAATCAGCATAATAACTTAGCAAGCTAATTATAAGGAGATGAAATTGGAATCGCCACTAGGTTCTGATCTGGCACGGTT | 9 | 53 | | 121 | | CBW17472 |
| sRNA1479529 | ACCCACTGATTATTTCTGATCAACGCCATGTCTGATAACGACCAATTGCAGCAGATCGCGCATCTGC | 75 | 120 | | 35 | | CBW17476 |
| sRNA1492264 | CAGCCTCCAATTCGTTGTGGCGGTGTTGTAGTCATATCTTCTCCGGGGTTACGCGCATTAACGGTCATGCCGCTCATTTT | 191 | 30 | | 12 | | CBW17488 |
| sRNA1525196 | GACGAAACATCGCGCGCAGATCCGGGTAAGCGTGTAAACGATGCATGAGCGGGTAAACAGGGAGTTTCAGGAT | 286 | 238 | | 99 | | CBW17515 |
| sRNA1529092 | GAGAAAAGGCAGGTAAAATCCTGCCTTTTGTGGGGATTAACGGAATAACGGTTTTACAGCGACGGGAAT | 69 | 30 | | 5 | | CBW17520 |
| sRNA1545620 | CAGGGATTACCCGGCCCGCAGACCGGGTATAACCCCTTAGCTACGGTGGACGCTGAGCCCTGCGAAGGATTGGGTAA | 259 | 17 | | 0 | | CBW17536 |
| sRNA1561940 | ATCAACTGACCTGAGCTCGACTTAACTGGCGCAGAATGTTCATCTGCGCCATAAC | 10 | 34 | | 10 | | - |
| sRNA1672712 | GCCCGCTGGCGCGACGCTTACTAAAGGAAGGATGATCGCATCGTCGCGCCAGTATCATGT | 11 | 381 | | 202 | | CBW17651 |
| sRNA1697847 | AGCCTGCCCGAACGCGCTGTTCTAATCCTGTCGCTCCCCTGCGCTCAGGGGAGCATTTTCGGATAACCCCTGAAGATCTATCCGGTCAT | 33 | 111 | | 12 | | - |
| sRNA1711201 | ATATCAATTACGGCTTGAGCAGACCTATGATCCCGGAAAAGCGAATTATACGGCGCATTCAGTCTGGCGGTTGTGCTAT | 143 | 7 | | 6 | | CBW17686 |
| sRNA1732405 | CGACGTTAACTATAAGTAAATAGGAACATAATTATGTCACAGACTGTACATTTCCAGGGTAACCCGG | 3 | 13 | | 0 | | CBW17707 |
| sRNA1804100 | AACAAACCACCCCAATATAAGTTTGAGATTACTACAATGAGCGAAGCACTTAAAATTCTGAACAACATCCGTACTCTTCGTGCGCAGGCAAGAGAAT | 0 | 108 | | 588 | | CBW17777 |
| sRNA1832521 | CAGGTGCCGTTTTATGGTTAATACCGAGCGCTAAAAGGGTCATGTCTGCGGGAGTAGTACCAGCGTTGATATGGTTAGTCT | 187 | 53 | | 24 | | CBW17800 |
| sRNA1837680 | GTTTTTGCGTTTCATATGTCACCTCCGGAACTTTTGGGTTGTTTTTAGGAAATCCCCCTAAACTTTAT | 29 | 110 | | 7 | | - |
| sRNA1900741 | AAGTGTTCATGCCAGGCCTGGCCTCCGTTTCAGAACACCAGGTGTTCTGCGCGTACAATCAAAGACATACCCGAAT | 64 | 13 | | 43 | | CBW17869 |
| sRNA1909049 | AACTGGCTGCGCCAGGGGAGGGTTATCCCCTGGCGCAGCGAGCTACTCACTATCGCCTGTTCACCTTCAATACTCTTCCTCCTTAAAAACATATCGGAAA | 66 | 117 | | 167 | | CBW17877 |
| sRNA1927927 | AATGGCTGTAGAGACTATGAAAAGGAAGTCATTATGTCCTCCTGGAAAATTGCTGCTGCGCAGTATGCGCCCCTGAAC | 53 | 6 | | 1 | | CBW17905 |
| sRNA1929359 | TTGCGGGCGCGCGTCTATGCGCTATGTAAGGTTCCCTGCGCCAGGCCGATTAAAAAG | 27 | 33 | | 2 | | CBW17907 |
|  |  |  |  | |  | | CBW19581 |
|  |  |  |  | |  | | CBW16226 |
| sRNA1978641 | AGGCAAAAAAAAGCAGCGGTAATGACTTACCGCTGCTGGAGTGTTTGTCCACACCGTTTCGGTTAAACAGCCTGTTCGATCTGTTCATCCAGCAGTTG | 158 | 92 | | 199 | | CBW17953 |
| sRNA2258692 | AGCTTTGAGATCGTCATGATGCTTTCCTCATCCTTTTACGGCCTTTTTCAGAG | 0 | 39 | | 73 | | CBW18239 |
| sRNA229689 | ATTTGCTACGTAAACAGAGTCGGTAAAAAAAAGCTGTGCCACATTTATGTCGGGTGGCAGCCATGCTCAACCCGACCTACA | 53 | 6 | | 0 | | CBW16297 |
| sRNA2408330 | ATCGGCGTCGTTCTGGTGCTTGCCAGTCTGCTCAGCGTTGGCGGTCAGCTCTGCCAGAAGCAAGCGACACGGCCATTGACGGCGGGCGGACGCCGCCGTC | 5 | 31 | | 10 | | CBW18373 |
|  |  |  |  | |  | | CBW19871 |
|  |  |  |  | |  | | CBW18321 |
| sRNA2425315 | TTTGGGTTTGTAGGCCGGATAAGCACTGCGCCATCCGGCAGTCGTTCATCCAGCCAACAAACTCAACGCAGTACCATCA | 276 | 204 | | 57 | | CBW20304 |
|  |  |  |  | |  | | CBW16297 |
|  |  |  |  | |  | | CBW18389 |
| sRNA2487522 | TGCCCAGGCCAGTAATCACTGCACGTTTCATTCAATACCTCTGTAATTCGCACTATTTTTAAGTTTC | 110 | 30 | | 11 | | CBW18449 |
| sRNA2513163 | CTCGCCCGTTGGGGGAAGCGTCACTCTACATCTGAAAAAGGCCGCAAGGCCTTTTATTT | 37 | 62 | | 3 | | CBW18471 |
| sRNA2514922 | ATACGCGTAAAGCGACGTTCAGGGCGGAAGTCAGCCATAATATCCTCAGATTAACGTTAGCGCCCGGACCGTCCGAGCGACGCTGCCACGATGGTGGCTC | 128 | 31 | | 42 | | CBW18472 |
| sRNA255259 | CTTTCGAGGATTTTAGAATGGCTGAAATAACCGCATCCCTGGTAAAAGAGCTGCG | 221 | 17 | | 2 | | CBW16320 |
|  |  |  |  | |  | | CBW18301 |
| sRNA2670362 | ATCCCGCCAGCGCCAGCATGATCATGCAGGCCACCACGCGTAAATGTTTCATCTTTCATCCCTGGCCTTAACGGCCTGTCAGCAACGGTTGAT | 367 | 179 | | 94 | | CBW18595 |
|  |  |  |  | |  | | CBW18432 |
| sRNA2681974 | AGAGCCCATATTGCCGGTGTGGGAGGTTTCCACCAGCACAATTCGAATGTTTTGCAGCATTGTCTTTCTTCAGCTAAAGATTAT | 13 | 16 | | 12 | | CBW18608 |
| sRNA2701330 | GGTGGTTCGCTGCCCACAAGGAGGAGACGGGCAGCGAATCGGCACATCAGATAGCCGCGTCGTCTTCTTCGCCAGTACGGATACGAAT | 96 | 16 | | 10 | | CBW18624 |
| sRNA2724415 | ATAAAATGCCCGTCACCAGTGTGGCTATCACCAGAATCAGGGCAAACATGTTCGCCATGCCAACTCCTTAGGGAT | 18 | 46 | | 3 | | CBW18644 |
|  |  |  |  | |  | | CBW20373 |
| sRNA2797098 | CTTTCGTTCCCGAAAATCCGGAATCACTGCTGAAAATCGTCTTCTCATCCGCCAAAACATCTTC | 12 | 23 | | 5 | | - |
| sRNA2837659 | TGGGTAACACACCCGTTACCGAGATCCGCCCGAAGATGCTCAAGGAACACCTCACCCCGCTGGAAGAACAGGGC | 42 | 109 | | 4 | | SL1344_2723 |
| sRNA2935765 | AAAAAGCCCCGAATTCACGGGGCTGAATAAAACGAAATAAATTAACGTAACAGAGACAGCACGTTCTGCGGGA | 625 | 54 | | 14 | | CBW18855 |
|  |  |  |  | |  | | SL1344_4369A |
| sRNA3075405 | CTGAAAAGTCTGGTGTAAACGTGTCAGACCCGATAGCGCCGCGCTATCGGGTCTGCGCG | 36 | 93 | | 39 | | CBW18987 |
| sRNA3089101 | TCCGCCGCTTTATCGCTGCGGTAAATAACGCAGCGGGTTTACGGATT | 9 | 2 | | 0 | | CBW19003 |
|  |  |  |  | |  | | CBW1750 |
| sRNA3120172 | GTGCACCGCTGAGGGTGCACCGCCGTTCTAACTTTTTTACTTCGCCTGACGTTTCTGATGCTCGTTGGCGGCTTTAACAAAGCCAGCGAACAACGGATGT | 789 | 2585 | | 994 | | CBW19031 |
| sRNA3124151 | GCAGTTCATCGCCAAGATCAACGGTCATTGTTCTGGCCATAACAGCCTCC | 190 | 12 | | 0 | | CBW19035 |
| sRNA3185694 | TTCTTTTTTTGGCAGGACTTTGTTCGATGTTATTTGCATGGATAACC | 149 | 1 | | 0 | | CBW19083 |
|  |  |  |  | |  | | CBW18735 |
| sRNA3226124 | AACGCTATAGCAGAATCGAGATGTAACGTAATGAACATGGGTCTTTTTTACGGTTCCAGCACGTGCTACACC | 8469 | 13359 | | 1012 | | CBW19120 |
| sRNA3271556 | CCCTGTTCGCCTGCTGACGAAGGCGAACCCATAGACATGTCGTCAGACATAGCGAACCTCAAT | 9 | 93 | | 7 | | CBW19566 |
|  |  |  |  | |  | | CBW19679 |
| sRNA3287406 | TAAAAAGGAAAGATTACTTCAGGCTGCCAACCAGCGCTTTGCGGCTATCTTCTAAGGTCACTACGCGGCTACAGACA | 47 | 32 | | 62 | | CBW19181 |
|  |  |  |  | |  | | CBW18253 |
| sRNA3373745 | ATTTTGAAGGTCAGGTTCGGACGGCAAATTGATAAGGAGTAATGGATGTCAGTACCACTTATTCTGACCTTACTGGCGGGCGCCGCCACCT | 106 | 98 | | 24 | | CBW19263 |
|  |  |  |  | |  | | CBW19535 |
| sRNA3391135 | AAAGAGAAGCGCTGCCCGAACCCCGCCAGCTAAACGCCATCCGGCACGATAAATTGCCTGATGGCGCTGCG | 32 | 81 | | 115 | | CBW19276 |
|  | CAGGGATTACCCGGCCCGCAGACCGGGTATAACCCCTTAGCTACGGTGGACGCTGAGCCCTGCGAAGGATTGGGTAA |  |  | |  | | CBW19535 |
| sRNA3397096 | ATACCGTCTTATGGAGCAAAACCCGCAGTCACAGCTGAAACTTCTTGTCACCCGTGGTAAGGAGCAAGGCTATCTGAC | 36 | 72 | | 35 | | CBW19283 |
| sRNA3399807 | AATTAAGCCACTCTAAAGAGTGGCCTTTTTGTTTCTGACAGGGCTCACCCCATACCGCTATTTACGATGCCAAT | 2 | 32 | | 56 | | CBW19285 |
| sRNA3411680 | AAAAATCCCTTTCTGGAGTCCGCTGACGTTTATGATACGTTTCGCTGTAATTGGGACTAACTGGATC | 46 | 30 | | 10 | | CBW19294 |
| sRNA3420128 | TTTCTGTCGGACCGTGGCAGGATCTGCTCGAACCTCACGGGCGCGACGTGCGCTTTGGCTAATTTCAGGCGTCCTCCGGGACGCCTTATT | 137 | 23 | | 2 | | CBW19303 |
| sRNA3474230 | CGTAAATCCCCGGGAGCATAGATAACTATGTGACCGGGGTGAGCGAGCGCAGCCAACGCCGTTGCGGCCCAAAAGACGACGAAGATTAA | 7 | 53 | | 6 | | CBW20496 |
| sRNA3536250 | ACGGAAACATAAGACAACGGGAGATGTTCATGACCTGGGAATATGCGCTAATTGG | 339 | 1482 | | 922 | | CBW19415 |
| sRNA3561197 | ACCTTGCTTTGCCGGACGGCGGCGTGAACGCCTTATCCGGCCTGCAAATACTCGTACGGGTCGCGGTCTTTTACA | 498 | 11 | | 1 | | CBW20304 |
|  |  |  |  | |  | | CBW18887 |
|  |  |  |  | |  | | CBW16297 |
| sRNA3576659 | CTTGCAGTCACAGTATGGTCATTTCTTAACTCATGCGCATCGGACAATATCAGCTCAGAAATCGCCTGATCGCAGCGCCCA | 355 | 516 | | 10 | | CBW19452 |
| sRNA3598379 | AATAATCTTTTCAACTCCTGAACACAACTCTGGATAATTATGTCAGTTTTGCAAGTGTTACATATTCCGGACGAGCGCCTTC | 284 | 11 | | 28 | | CBW19468 |
| sRNA3626080 | AAAAAACACTCCTTTTCAGGAGCCTGTCGTTATGTTTTCAGGGCAGGCTCATTATTTACGCGG | 1505 | 233 | | 7 | | - |
| sRNA3644974 | AATAGAAAAGAAATCGAGGCAAAAATGAGCAAAGTCAGACTCGCTATTATCGGTAATGGTATGGTCGGCCACCGCTTTATTGAGGATCTCCTT | 64 | 1 | | 0 | | CBW19536 |
| sRNA3710010 | AACGAACATTTTTGAACTTTAACGAAAGTGCAAGAGGGCAGCATGGAAACCAAAGATCTGATCGTGATAGGCGGGGGCATTAACGGTGCAGGCATCGCGG | 22 | 15 | | 53 | | CBW19587 |
| sRNA3763008 | AGAGCCGCAAGCATACCGTTTGCGAAAAAATAACGAATTCAAGGAATTAAGATGCTTTGGTCGTTTATCGCTGTCTGCCTGTCC | 29 | 16 | | 0 | | CBW19634 |
| sRNA3766133 | ACCGTCTACAAAATAAGGAATGTTAGGCACTATGAAAAATATCAAAGTCATCACCGGCGTTATCGCGACGCTGGGCATATTTAGC | 44 | 14 | | 1 | | CBW19636 |
| sRNA3770708 | TACGCAGATTGAGGCGCAATCCGTTTAGCGCCGGTTCAGCGACGGGTTCGGGCATGAGTATCACCGCATTTTTACAACAT | 44 | 14 | | 1 | | CBW19640 |
|  |  |  |  | |  | | CBW16141 |
|  |  |  |  | |  | | CBW18541 |
| sRNA3808647 | GCGTAGCTGCTGGATAACCTGCTTTATCATCTCAGTTGTCCTGTTAATGGGAGTGTGACTGTCCGGACTCGT | 59 | 119 | | 170 | | CBW19670 |
| sRNA3890778 | ACCCGTATTCACCTGGCTGAATACGTGTTGAAGGGGGATTGTTCGTCATGACCAACTCCTTATTGTAAAGGTAC | 102 | 28 | | 5 | | CBW19738 |
| sRNA4019064 | ACCACTGCGCGAGATGGCGCAGTTGTTAGTAGGGTCGCGTTCAGCATGGAAGGGTTCATTAGGGGCCTTGTCTGTTTGTTATTAAC | 243 | 34 | | 0 | | CBW19854 |
| sRNA4110510 | GCTGCTCAGACGGGAAATTCTTTCCGCTAATAAATGTGGGTGAGCCATAATAAAGAGGCGTCCTTTCGACAAATGGATCATGCT | 67 | 39 | | 5 | | CBW19937 |
| sRNA4150758 | AACCCGTCGCCGCACGATTTAGCAACACCTCTGCAG | 16 | 8 | | 1 | | CBW19968 |
| sRNA4173538 | AGAAAGCATGAACTTACAGGCAATCCATAATGAAAAACGTGTTTAAGACACTCGCTGTACTTCTGACTCTGTTCAGCCTGACGGGCTGCGGTTTGAAAGG | 37 | 21 | | 24 | | CBW19991 |
| sRNA4300353 | ATCATTTCGATTTGGCAGAACTGGCCAGGCAAATCGCAGGGTAGCGACTTCACCCTGCCGGGGACGGTAAAACGTCCCCTGCTGGCGAAGA | 35 | 4 | | 0 | | CBW20105 |
| sRNA4303605 | CCTGATTCAGTATGGGTACAGAGTCGAGCCATTTTTTTATCCTCGGCTATTTTTACATCGTTACATAAAAT | 93 | 28 | | 4 | | CBW20110 |
| sRNA4304655 | TAATCAGATACCAACGTATTATCGCTCATTGTCATAACCTGGCTTTACTTTGAACATTTCTAAATCATT | 124 | 13 | | 0 | | CBW20111 |
| sRNA4327130 | TTACTATTAACTTCGCGTGGTGTCTGGCGTTAGGGCTGGAAGAGCGACGCGGCCTTACACTGAGGTTCCCCATGAAAAAAGGTATTCACCCGA | 78 | 92 | | 41 | | CBW20134 |
|  |  |  |  | |  | | CBW18380 |
| sRNA4330526 | TAAAAAAGCGCGGCCAGAGGCGTTCTGACCGCATGCTTTGCTATCAGTATTCCCACGTTTCCGGATCAATCCCCAACTCACGCATGAT | 52 | 8 | | 2 | | CBW20137 |
|  |  |  |  | |  | | CBW20392 |
| sRNA4356426 | AGCCCAGGGCTTTCCAGCTAAACGCGGGTTTAGTCTGGAGTAATGTGGATTGCATAATGAGAGCCTGTCTTTAATCGTA | 21 | 5 | | 1 | | CBW20156 |
| sRNA4435181 | ATCAGGACGAGGAACGGGGCGAGTATTAAGTGAAATGCCGGATGGCGGGGCCTACGGAGAACATGCGCGTAGGCCCGTCATTCGGCATTCAATA | 47 | 16 | | 3 | | CBW20212 |
|  |  |  |  | |  | | CBW16297 |
| sRNA4475966 | AGGTTATGAATCGGAGAGTAAGGCATGTCACACCCTGCGTTAACGCAACTGCGTGCGCTGCGCTATTTT | 147 | 54 | | 16 | | CBW20257 |
| sRNA4490563 | TCCTGCCCTCTGTAAACCTGGAGAACCATCGCGTGTTTCAAAAAGTTGACGCCTATGCCGGCGATCCGATTCTTTCACTGATGGAGCGTTTTAAAGACGA | 1617 | 17 | | 1 | | CBW20271 |
| sRNA4520408 | TTACGCTACCTGTAGATAATCAAGAAGAACACGCATAATAACGGAGGCCCCTCACCTTTGGGTGAGGGGGTTTACTT | 66 | | 5 | | 0 | CBW20284 |
| sRNA4550869 | AAGGGGGAGGGCGAGCCTCCCGGGGAGTTAGGTGGCAGGCTGATGTTTAA | 72 | 2 | | 1 | | CBW20311 |
| sRNA4614185 | AAGGGGACGATTCTAACGACGCCTAGCGTATCTGTCAGTCGTCCGTATCAGAAA | 36 | 5 | | 3 | | CBW20372 |
|  |  |  |  | |  | | CBW16754 |
| sRNA46165 | ATTCACCTGATTGAGAGAATAAAAAGTGAAACATCTGCACCGATTTTTTAGCAGCGATGCCTCTGGGGGCAT | 13 | 80 | | 54 | | CBW16141 |
| sRNA4641836 | GCCATCCGGCGCGAGCCACGCGGCATCAGGCATAGAGTATCGCCTGTACGCGCCAGTTATCCGACTGTTGTTCTTCATACATTTGAATAAT | 34 | 6 | | 21 | | CBW20397 |
|  |  |  |  | |  | | CBW20483 |
| sRNA4642515 | GCACGCTACACGCGCTTAATAACAGCGCTGCTATCAATGAAGCAAACCGACGCATAACTCTATCTCGTTTCCTGAT | 491 | 28 | | 24 | | CBW20398 |
| sRNA4655124 | ATTCTGTAAGCGCCCCGGGTTTGCCGGATGATGGCGTAAACGCCTTATCCGGCAACCCTGGCTTCA | 73 | 25 | | 12 | | CBW20304 |
|  |  |  |  | |  | | CBW18887 |
| sRNA4672451 | ATTGAAAGCGGTGACTGGTTAGACAGAAACCAGCCCTGAACTGAACACGCCACCCTCGGGTGGCGTTGTTTTT | 43 | 13 | | 2 | | CBW20430 |
| sRNA4723568 | CCCGAACCGGGCAAACTTGCGCCGCCCGGCGGGGATGACGTCCGGCGCTTAGCGACGAACAGCGAT | 59 | 1 | | 0 | | CBW20477 |
| sRNA470505 | ACCCTGGAAGAGAGAATATGAACATTATTAAAGCTAACGTTGCAGCCCCTGACGCTCGTGTCGCCATCACTATTGCGCGTTTCA | 59 | 49 | | 14 | | CBW16511 |
| sRNA4784864 | TTGATAGTGTGTTTCTGACCTTGACGTAAATAAGCGGGAAAAGCGATGAAACACACCATTGGGATACTGGGGGGGATGGGACCTGCGG | 41 | 18 | | 2 | | CBW20529 |
| sRNA4810641 | ATCTGTCTAGCGGAAAGAGAAAACATGTTAAAGCGAATTAAAATTGTTACCAGCTTACTGCTGGTATTGGCGCTATTT | 32 | 72 | | 184 | | CBW20553 |
| sRNA663093 | CGCTATGGTGGAAAAGGCGGCGCGCGTTGCTGCTGAGGGTTGCCAGGCCTGATAAACGCAACGTCACCAGGCTATATT | 143 | 32 | | 9 | | CBW16689 |
|  |  |  |  | |  | | CBW16689 |
| sRNA692584 | CTTCCGCTGCAAACGTAACTGCTCTGTAAGCATACGTCCGACAGCAAGA | 38 | 63 | | 16 | | CBW16717 |
| sRNA732133 | AATGTAGGCCGGATAAGGCGCAAGCGCCGCCATCCGGCAAGCTACTATTCTGCAATCTT | 62 | 19 | | 3 | | CBW20304 |
|  |  |  |  | |  | | CBW16297 |
|  |  |  |  | |  | | CBW19985 |
|  |  |  |  | |  | | CBW18887 |
|  |  |  |  | |  | | CBW17016 |
| sRNA733846 | AGGAAGCCCGACAGGCGTAGCGCCATCGGGCATTATTGCCTTTTCAGTAGGGTTAAGGCTGGTAGAATCCTACGCCCAGCTCATTTTCTTTA | 87 | 34 | | 12 | | CBW16758 |
| sRNA746802 | ATACGTTGTTTACGCTTTGAGGAATCCACGATGAGTGAGGCTGAAGCCCGCCCGACTAACTTTATTCGT | 38 | 34 | | 18 | | CBW16767 |
| sRNA7619 | AGACCGGTTACATCCCCTAACAAGCTGTTTAAAGAGAAACTCTATCATGACGGACAAATTGACCTCCCTTCGTCAGTTCACCACCGTAG | 301 | 172 | | 105 | | CBW16108 |
| sRNA815338 | ATTATCGTGGGTCGCAGGCCCAGGTATGGGAGATATGATGAAGCAGGCATTACGAGTAGCATTTGGTTT | 63 | 37 | | 26 | | CBW16828 |
| sRNA836354 | GCTTAGTTACAGCCATACTCTCACTCCTATAAATCATATGTAATGATAATAAT | 107 | 130 | | 214 | | CBW16847 |
| sRNA924314 | GAACCCTGGTAGAAAAGAGGAGGAAAAAAGGTATGGTCGTTGACAGACTGAGAACCGATCTTCTCAACAAGCTGATAAACGC | 34 | 45 | | 25 | | CBW16926 |
| sRNA944582 | AAAATAACAGTAAACATATCTCTCCTCGAACATAAAGCCAGACGGCGAAC | 5 | 13 | | 15 | | CBW16945 |
| sRNA957893 | AGGCGAGTAATGCAGCCAGAACTAACTTTTTCATGATGTCACTCCCGTAATCTTATGAT | 113 | 51 | | 10 | | CBW16959 |

^a^ Log – ; 5h – ; 24 –

^b^ Putative Gene Targets –
